# Genomic introgression between critically endangered and stable species of Darwin’s tree finches on the Galapagos Islands

**DOI:** 10.1101/2024.05.30.596739

**Authors:** Rachael Y. Dudaniec, Sonu Yadav, Julian Catchen, Sonia Kleindorfer

**Author notes:** Corresponding Author Rachael Y. Dudaniec, School of Natural Sciences, Macquarie University, Sydney, 2109, Australia. present address (Sonia Kleindorfer).

## Abstract

Natural hybridisation among rare or endangered species and stable congenerics is increasingly topical for the conservation of species-level diversity under anthropogenic impacts. Evidence for beneficial genes being introgressed into or selected for in hybrids raises concurrent questions about its evolutionary significance. In Darwin’s tree finches on the island of Floreana (Galapagos Islands, Ecuador), the Critically Endangered medium tree finch (C*amarhynchus pauper*) undergoes introgression with the stable small tree finch (*Camarhynchus parvulus*), and hybrids regularly backcross with *C. parvulus.* Earlier studies in 2005-2013 documented an increase in the frequency of *Camarhynchus* hybridisation on Floreana using field-based and microsatellite data. With single nucleotide polymorphism (SNP) data from the same Floreana tree finches sampled in 2005 and 2013 (n = 95), we examine genome-wide divergence across parental and hybrid birds and evidence for selection in hybrids. In assessing previous estimates of introgression we found that just 18% of previously assigned hybrid birds based on microsatellites were assigned to hybrids using SNPs. Over half of the previously assigned hybrids (63%) were reassigned to *C. parvulus,* though parental species showed concordance with prior assignments. Of 4869 private alleles found in hybrid birds, 348 were at a high frequency (≥0.30) that exceeded their parental species of origin 89-96% of the time. Across the two years, 3436 (70.6%) private alleles underwent a substantial (≥0.30) allele frequency increase or decrease. Of these, 28 private alleles were identified as candidate loci under selection via local PCA genome scans and outlier tests.

Alleles were annotated to genes associated with inflammation, immunity, brain function and development. We provide evidence that introgression among a critically endangered and stable species of Darwins’ tree finch is being retained by selection across years and may aid in the retention of genetic diversity in birds threatened with extinction.

## Introduction

Natural hybridization and introgression occur in at least 10% of animal species (Mallet 2005) and can play a significant role in increasing genetic diversity or creating novel genotypes that affect hybrid fitness (Bar-Zvi et al. 2017). Despite its potential for incompatibilities and deleterious fitness consequences, hybridization may allow individuals to more quickly occupy new ecological niches or embark on novel evolutionary trajectories (Kalumni et al. 2024; Abbott et al. 2016). Although adaptive introgression appears to be common (Mallet et al. 2016) and most introgressed variation is selected against (Martin and Jiggins 2017), preferential introgression of locally adapted alleles into hybrids may facilitate increased fitness (e.g., Patton et al. 2022; Arnold et al. 2016; Fraisse et al. 2014). Indeed, when compared to parental populations, hybrids may exhibit a fitness benefit when challenged with novel environmental conditions, for example, during range expansion (Stahlke et al. 2022), invasion (Qiao et al. 2019; Hovick and Whitney 2014), pathogen exposure (Peters et al. 2019; Fritz et al. 1999) or anthropogenic impacts (Thaweepworadej et al. 2023; Ottenburghs 2021). Such challenges for biodiversity are ubiquitous under current climate and land use change, and hybridisation has been proposed as either a natural or managed mitigation strategy for the conservation of some susceptible species (Kalmuni et al. 2024; Chan et al. 2019).

Although hybridisation can have beneficial outcomes for species or populations via adaptive introgression (Kulmuni et al. 2024), it may also lead to reduced biodiversity via species collapse (i.e., fusion of two distinct species or lineages into one), genetic ‘swamping’ (e.g., reduced fitness due to maladaptive hybrids) or even extinction (Arce-Valdés and Sánchez-Guillén 2022, Bog et al. 2017, Kleindorfer et al. 2014). Therefore, the underlying genetic characteristics and fitness outcomes of hybridisation require study to ensure appropriate management of biodiversity within specific hybridisation contexts.

Within adaptive radiations on islands, introgression among closely related species can be important for the evolution of novel fitness peaks (Patton et al. 2022). Species that have undergone relatively recent adaptive radiations often exhibit weak genetic divergence, with phylogenetic lineages best separated with fast evolving markers such as microsatellites (Takezaki et al. 1996) or single nucleotide polymorphisms (SNPs) (Leaché and Oaks 2017), as opposed to more conserved genomic regions (e.g., mtDNA: Ballard and Rand 2005). This is particularly relevant for Darwin’s finches on the Galapagos Islands in Ecuador, which have reused ancestral modules of genetic variation for beak size over the past million years with introgression, leading to the rapid speciation (Rubin et al. 2022). Further, Darwin’s finches are a relatively recent radiation of species, such that previous efforts to untangle the species diversity of finches across islands have employed microsatellites (Petren et al. 1999), or whole genome phylogenetic analyses (Lamichhaney et al. 2015). While microsatellites are too sparse and whole genome analyses too impractical in large numbers of individuals, ascertaining the incidence of hybridisation in closely related species has been enabled by the generation of dense SNP data sets via high-throughput sequencing technologies.

Compared to other species groups, the reduced genetic divergence among Darwin’s finches has led some to conclude that standard evolutionary paradigms cannot be widely applied to this complex adaptive radiation (Zink and Vázquez-Miranda 2019). Such challenges are similar for the identification of hybrid birds within the group, given the relatively recent divergence of the Darwin’s finch assemblage over the past 1.5 million years (Grant and Grant 2014). Hybridisation and bi-directional introgression among Darwin’s finches (*Geospiza* spp.) have been well documented on the island Daphne Major, with findings of rare (1-2%) but consistent rates of hybridisation across years that are affected by rainfall and resource availability (Grant and Grant 2008, 2014, 2016).

Floreana Island on the Galapagos Islands contains two species of Darwin’s tree finches (*Camarhynchus* spp.) that are undergoing hybridisation, the small tree finch (*C. parvulus*) and the IUCN-listed Critically Endangered medium tree finch (*C. pauper*) (Peters et al. 2017, Kleindorfer et al. 2014). The common occurrence of hybridisation documented in this population co-occurred with the confirmed absence of the large tree finch (*C. psittacula*) and hence could have been a result of species collapse (Kleindorfer et al., 2014; Peters et al. 2017). In 2005, the large tree finch was considered rare and possibly absent on Floreana Island (Grant et al. 2005), and its presence could no longer be confirmed in 2010 (Kleindorfer et al., 2014). Instead, there was evidence for one-way genetic introgression from the extant larger-bodied finches (*C. pauper*) to smaller bodied finches (*C. parvulus*), that was largely driven by *C. pauper* females pairing with *C. parvulus* males (Peters et al., 2017, 2019). The resulting hybrid birds had intermediate body size, and their song was indistinguishable from that of the male parental species (*C. parvulus*) (Kleindorfer et al. 2019; Peters & Kleindorfer, 2018). Finally, Kleindorfer et al. (2014) used microsatellite data and found a 73% increase in the percentage of *Camarhynchus* hybrids between 2005 and 2010 following a period of drought and then high rainfall (Kleindorfer and Dudaniec 2020).

Another potential driver of this hybridisation are impacts from the introduced myiasis- causing ‘avain vampire fly’ *Philornis downsi* (Common, et al. 2019; Fessl et al. 2006), which causes anaemia, mortality and deformations of the nestling naris via larval blood and tissue parasitism (Common et al 2023, Katsis et al. 2021, Kleindorfer and Dudaniec 2016). The number of *P. downsi* larvae in the nest predicts the size of naris deformation (Katsis et al. 2021) and adult finches with enlarged nares from early-life parasitism produce low quality song that no longer differs between small and medium tree finches (Kleindorfer et al. 2019). By this mechanism, species recognition may become blurred, and parasitism may promote hybridisation. Notably, hybrid tree finch males on Floreana island had 50% fewer *P. downsi* parasites in their nests compared with *C. parvulus* nests and 60% fewer parasites than *C. pauper* nests (Peters et al., 2019). The effect was also evident according to the inferred degree of *Camarhynchus* hybridisation, with increased genetic admixture of the attending male correlating with decreased *P. downsi* within nests (Peters et al. 2019).

This study examines the significance of hybridisation for adaptation by asking: which genes introgress across species boundaries, and among those genes, which ones show evidence of selection? Given the temporally shifting observation of hybridisation frequency in the Floreana tree finches (Kleindorfer et al. 2014; Peters et al. 2017, Kleindorfer and Dudaniec 2020), this study system offers an opportunity to examine the genomic characteristics of introgression, particularly for loci that may be uniquely selected for advantageous hybrid traits. Such observations may be relevant for understanding the early mechanisms of speciation and evolutionary responses to climate change and parasitism. With a comparison of the Floreana Island Darwin’s tree finch population sampled in 2005 and 2013, we aim to: 1) verify previous hybrid genetic assignments based on microsatellite analyses with a new, high resolution genome-wide SNP dataset, 2) examine genome-wide divergence across the parental and hybrid tree finches, and 3) establish whether hybrid tree finches inherit private alleles that are unique to each parental species, and whether those alleles show evidence for selection or map to gene functions relevant for hybrid fitness.

## Methods

### Species sampling and study site

The study site was in the highlands of Floreana Island (01 17°S, 090 27°W, 300-400 m asl), Galapagos Archipelago, within its humid *Scalesia* forest at the base of Cerro Pajas volcano. Our study was conducted during the finch breeding season from February to April of 2005 and 2013. The focal tree finch species were small tree finch (*C. parvulus*), medium tree finch (*C. pauper*), and the recently discovered hybrid group that arises from pairings between *C. pauper* females and *C. parvulus* males (Kleindorfer et al., 2014; Peters et al., 2017), which all occur in sympatry in the highland habitat dominated by *Scalesia pedunculata* forest at 400-600 m elevation.

Darwin’s finches were captured in 6 x 12m mist-nets between 600-1100 hrs. Each bird was banded with a numbered aluminium band and a unique combination of colour bands.

Morphological measurements and blood samples were taken from adult tree finches as described in Dudaniec et al. (2006) and Kleindorfer et al. (2014). Blood samples (10ul) were immediately transferred to Whatman Classic FTA paper for DNA preservation. All protocols and procedures employed were ethically reviewed and approved by the Flinders University Animal Welfare Committee under permit numbers E190 and E270.

### DNA extraction and RAD sequencing

A total of 95 samples of adult tree finches (*Camarhynchus* spp.) were selected (2005: N = 56; 2013: N = 39) from samples that were previously identified as parental or hybrid birds using genetic assignment analyses with 10 microsatellite markers (Kleindorfer et al. 2014; Peters et al. 2017). These studies implemented assignment cut-offs of 0.75 (*C. pauper*) and 0.80 (*C. parvulus*) for parental species assignment using the program STRUCTURE (Pritchard et al. 2000). Based on these prior microsatellite assignments, we extracted DNA from the following individuals. In 2005: 13 *C. parvulus*, 22 *C. pauper*, and 21 hybrids. In 2013: 12 *C. parvulus*, 10 *C. pauper*, and 17 hybrids. We extracted DNA from Whatman Classic FTA paper using a modification (200 lL volumes used for all washes) of method 4 from Smith & Burgoyne (2004). Two extractions were undertaken per sample, which were then combined, concentrated via evaporation, and quantified on a Qubit 2.0 Fluorometer to obtain 20-40 µl DNA per sample. We constructed Restriction site-Associated DNA sequencing (RADseq) libraries from the samples using the SbfI restriction enzyme, similar to the method of Baird et al. (2008) and following a protocol performed by Floragenex, Inc. (Oregon, USA), as described in Supplementary Text S1. Samples were paired-end sequenced on the Illumina HiSeq4000 platform.

### Genetic marker characterisation

Raw sequences from each RAD library were quality checked using FASTQC (Andrews et al. 2010). Each library was demultiplexed using *Stacks v2.54* (Rochette et al. 2019; Catchen et al. 2011, 2013) and reads aligned to the *Camarhynchus parvulus* reference genome STF-HiC (1.28 GB; Genbank accession number: GCF_901933205.1) and loci were assembled with gstacks as described in Supplementary Text S2 and Table S1. The populations program within *Stacks* was used to filter SNP loci such that SNPs within a RAD locus had a minimum minor allele frequency (MAF) of 0.03 and were found in a minimum of 75% of individuals within a population. The populations program was run in two stages, 1) with populations defined according to the two parental species and hybrid groups identified from previous microsatellite analyses (Kleindorfer et al. 2014; Peters et al. 2017), and 2) following neutral genetic structure analyses (using the dataset from stage 1), with populations defined according to the revised SNP-based genetic cluster assignments to either parental species or hybrid group.

After stage 1 filtering, there was a total dataset of 10381 SNPs. A second run of populations was performed with identical parameters but specifying the --write-single-snp flag to retain a single SNP per RAD locus, thus excluding SNPs that are physically linked. This resulted in a SNP dataset of 6637 SNPs for analysis of neutral genetic structure. Individual missingness (i.e., the percentage of missing data per individual specimen) was calculated in *vcftools* v0.1.13 (Danecek et al., 2011). For stage 2 filtering (i.e., after undertaking genetic structure analysis – see below), we repeated the above, resulting in 10463 total SNPs. When using the --write-single-snp flag 6757 SNPs were retained. The stage 2 dataset was used to examine private alleles and genetic divergence within and between each genetic cluster (see below).

### Genetic structure and hybrid re-identification

Using 6637 SNPs (from stage 1 filtering), we first assessed pairwise genetic relatedness between our samples to eliminate any full sibs that may be in our dataset. Pairwise relatedness was calculated using the method of Manichaikul et al. (2010) implemented in *vcftools* v0.1.13 (Danecek et al. 2011). We assessed the microsatellite-based species and hybrid assignments of our samples (Kleindorfer et al. 2014; Peters et al. 2017) by conducting genetic structure analyses using our 6637 SNP dataset. First, a principal component analysis (PCA) was performed using *pcadapt* in R (Luu et al. 2017). The software *snmf* within the *LEA* package in R (Frichot et al. 2014, 2015) was then used to assign individuals to parental or hybrid groups using individual ancestry coefficients obtained from a sparse non-negative matrix factorisation (*snmf*) algorithm. The *snmf* program was run from K 1-10 with 100 repeats per K, iterations were set to 1,000 and regularization parameter α was set to 100. Optimal K was selected based on the lowest cross-entropy across all runs. Assignment probabilities (i.e. qmatrix values) were used to group individuals into parental (≥0.80 q value) or those having hybrid ancestry (<0.80 q value), in accordance with previous ancestry coefficient thresholds applied to these samples using microsatellites (e.g., Kleindorfer et al. 2014).

Samples were reassigned as being either hybrid or belonging to the *C. parvulus* or *C. pauper* species, according to the results of *snmf*, prior to subsequent analyses. Notably, we use the term *hybrid* to describe individuals that show shared ancestry with either parental species based on our assignment threshold (<0.80 q value), however we were unable to ascertain the number of generations of hybridisation within each sample. Thus, we expected a gradient of assignments to the hybrid group, given previous observations of a hybrid swarm in this system, which likely involves substantial backcrossing between female *C. pauper* and male *C. parvulus* or hybrid birds (Peters et al. 2017). For visualisation, a second PCA was further performed in *pcadapt* with reclassified samples. Following this, stage 2 filtering in populations was performed, as described above.

Pairwise FST was calculated between the revised genetic groups using the WC84 estimate implemented in the *assigner* R package (Gosselin et al. 2016). Bootstrapped 95% confidence intervals (CIs) were calculated using 100 iterations. Observed and expected heterozygosity and FIS were calculated for each genetic group (after stage 2) in *Stacks*.

To further examine patterns of genetic co-ancestry among our classified hybrid individuals and parental species, the updated, re-classified datafile based on *snmf* assignments was analysed in *fineRADstructure* in R (6637 SNPs; Malinsky et al. 2018). This program uses a Markov chain Monte Carlo (MCMC) clustering algorithm to infer population structure and is specifically designed to infer population structure amongst genotypes obtained from RADseq data. First, RAD loci were reordered based on linkage disequilibrium using the script sampleLD, provided within *fineRADstructure*, followed by calculation of the co-ancestry matrix with *RADpainter*. We then ran *fineRADstructure* with 100,000 burn-in iterations, followed by 100,000 sample iterations and 1000 iterations for file thinning. *FineRADstructure* uses haplotype linkage information and focuses on the most recent coalescence (common ancestry) among the sampled individuals to derive a “co-ancestry matrix” – a summary of nearest neighbour haplotype relationships in the dataset.

### SNP outlier analysis

We tested for FST outliers using our stage 2 dataset of SNPs (N = 10381; with individuals re- classified as parentals and hybrids) using three approaches implemented in 1) *Bayescan* (Foll and Gaggiotti 2008), 2) *OutFlank* (Whitlock and Lotterhos 2015), and 3) *pcadapt* (Luu et al. 2017). *Bayescan* uses a Bayesian framework to detect outliers with high FST while accounting for sample size (Foll and Gaggiotti, 2008), demographic processes and hierarchical genetic structure (de Villemereuil et al., 2014). Bayescan was run with 50,000 burn-in iterations, a thinning interval of 10, and sample size of 5000, resulting in 100,000 total iterations. The number of pilot runs was set at 20 with a length of 5000 for each run. We set the prior odds to 10 and implemented a false discovery rate (FDR) threshold of 0.05. Log-likelihood traces were plotted in R to confirm model convergence. *OutFlank* is based on the Lewontin and Krakauer (1973) method, while accounting for sampling error and nonindependence between sampled populations. *OutFlank* detects outliers under divergent selection by initially inferring the FST distribution from multiple loci and then fitting a χ2 model to the centre of the distribution, resulting in a null distribution. This null distribution is then used to detect outlier loci. We used a left and right trim value of 0.05 as suggested by Whitlock and Lotterhos (2015) with *K* defined by genetic structure analysis described above. Loci with an expected heterozygosity of <10% were excluded as recommended by Whitlock and Lotterhos (2015), and the FDR was set to 5%. The software *pcadapt* performs genome scans to detect genes under selection and is effective at handling admixed individuals while not requiring grouping of individuals into populations.

*Pcadapt* uses principal components analysis to account for population structure (Luu et al. 2017), and computes p-values to identify outlier loci based on the correlation between SNPs and principal components axis (PCs). The numbers of PCs are decided based on a scree plot, which displays the percentage of variance explained by each PC. P-values were adjusted based on the Bonferroni correction with an expected false discovery rate lower than 5%.

### Identifying private parental alleles and divergence in hybrids

Using the stage 2 SNPs (N = 10381), we calculated per SNP pairwise FST and ΦST in *Stacks* (Catchen et al. 2013; Rochette et al. 2019) for all loci in both *C. parvulus* and *C. pauper* and visualised this across each chromosome of the *C. parvulus* genome to search for peaks of genetic divergence between parental species (Figure S7). We identified private alleles within each of the parental species (as described in Text S1) and mapped them along each of the 30 chromosomes of the *C. parvulus* genome using the stacks-private-alleles program. We further examined the parental private alleles found in the hybrids along each *C. parvulus* chromosome, according to its frequency in the hybrid samples (N=15), comparing frequencies in 2005 and 2013 for both *C. parvulus* and *C. pauper* private alleles (Figure S7). To identify private alleles that showed increased genetic divergence in the hybrid group, we further examined alleles with a frequency ≥0.30, which equated to approximately 30% above the mean private allele frequency observed in hybrids. This was examined as a conservative measure of the most confident allele frequency changes in the dataset.

### Localized heterogeneity across the genome

To examine for shifts in localized heterogeneity along chromosomes, which may indicate selection, we used the program *lostruct v.0.9* (Li and Ralph, 2019). Local changes in genetic relatedness across the genome were assessed across each pair of windows using 100bp windows and 2 PCs. We visualised the resulting dissimilarity matrix across windows using multidimensional scaling (MDS) and variation along the genome was identified by choosing three extreme windows in the MDS plot. We then highlighted these window positions along the *C. parvulus* genome and created aggregated PCA plots from the extracted corners (Li and Ralph 2019). Start and end base pair information of each MDS outlier window was extracted.

### Combined evidence for hybrid selection

We integrated the outcomes of the above analyses (i.e., FST outlier tests, local PCA, private alleles, allele frequencies, and changes across years) to find loci that show evidence of selection in hybrids. Specifically, we mapped the 2005 and 2013 allele frequencies of all hybrid private alleles on to the *C. parvulus* chromosome-level reference. We subsequently mapped *pcadapt* outlier loci, and outlier windows from local PCA to identify private alleles or gene regions that correspond with putative outliers and change in frequency across the two sampled years. Those alleles with multiple lines of evidence for selection were identified as the best candidates.

### Gene annotation

RAD loci (∼400-600bp in length after assembling paired-end contigs in *Stacks*) that contained 100bp outlier windows from the local PCA and that contained hybrid private alleles with an allele frequency ≥0.3 were identified for gene annotation. Loci containing SNPs identified as outliers via *pcadapt* were also selected for annotation. These sequences were considered to be candidates under selection and were annotated to the *C. parvulus* genome using NCBI BLAST with the general nucleotide collection database (Johnson et al., 2008). E-value threshold was set to 0.05 and BLASTn was used to match highly similar sequences to the *C. parvulus* genome.

## Results

### Genetic structure and assignment of hybrid ancestry

None of the samples had missing data >71% for both the ‘write-single-snp’ and the ‘all SNPs’ dataset, thus all 95 samples were included in the analysis (mean missingness = 16%). Pairwise genetic relatedness was low across all samples (mean = −0.02 ± 0.0007 s.e.) and no full siblings were detected (Figure S1). Our initial PCA showed two genetic clusters in ordination space, with microsatellite-analysed hybrid individuals spanning both clusters, with the majority in the *C. parvulus* cluster (Figure 1a, Figure S2). Two individuals appeared as outliers, yet these samples did not have high missing data or high genetic relatedness and were therefore retained in the analysis (Figure 1a). The genetic clustering approach of *snmf* (i.e., with parental species and hybrids defined from previous microsatellite studies; Kleindorfer et al. 2014, Peters et al. 2017) using an 0.80 admixture coefficient (q-value) cut-off value indicated two genetic clusters and a third admixed group (K= 2; Figure 1c, Figure S3).

**Figure 1.**
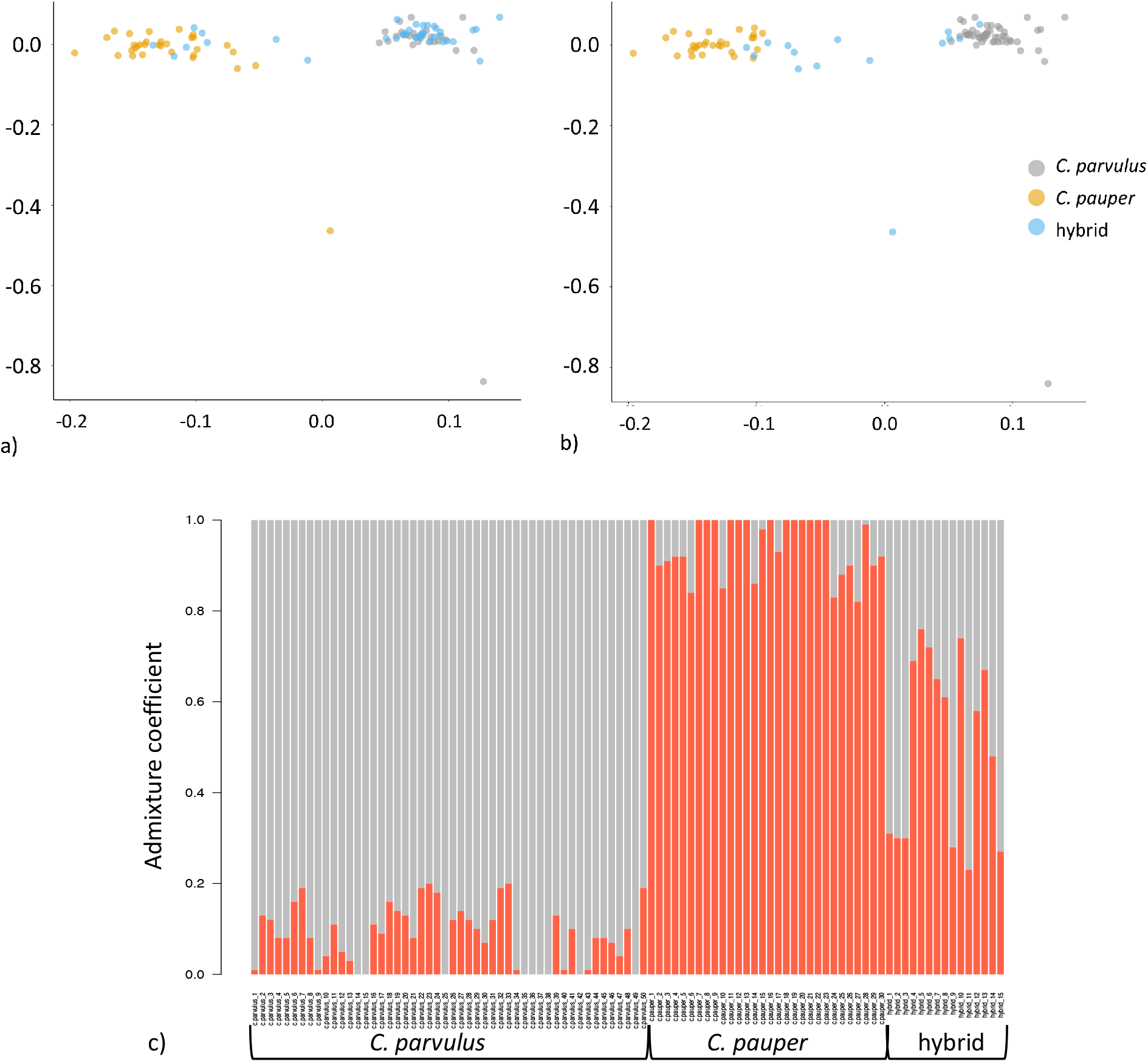
a) PCA plot from *pcadapt* with genetic groups labelled according to microsatellite-based genetic assignments; b) PCA plot from *pcadapt* with genetic groups labelled following SNP-based genetic assignment analysis (using *snmf*); c) Admixture coefficients (q-value) from genetic structure analysis using *snmf* showing two distinct genetic clusters, *C. parvulus* and *C. pauper*, with a third group with increased admixture (hybrid). Samples in c) are shown re- labelled after comparison with initial microsatellite identifications.

There were many discrepancies between the prior microsatellite-based classifications and the present analysis (Figure 1b). Of the 38 birds identified to have hybrid ancestry via microsatellite analysis, just seven birds (18%) were assigned less than the 0.80 q-value threshold for hybrid assignment using *snmf,* with the majority being assigned to *C. parvulus* (N = 24; 63%) and a few to C*. pauper* (N = 4; 11%). This large difference in hybrid assignments was not reflected in the microsatellite-identified parental species, with 26 of 28 birds (93%) originally assigned to *C. parvulus* being supported by our SNP data, and the remaining two given newly assigned hybrid status. Similarly, 26 of 32 birds (81%) originally assigned to *C. pauper* were supported by our SNP data, with six (19%) reassigned hybrid status. This resulted in a revised total of 50 *C. parvulus* (2005 = 24; 2013 = 26), 30 *C. pauper* (2005 = 21; 2013 = 9) and 15 hybrid birds (2005 =11; 2013 = 4) in our dataset, and all samples were re-classified accordingly (Figure 2, Table S1). The mean admixture coefficient (q-value) for birds in 1) the *C. parvulus* group was 0.915 ± 0.009 s.e, 2) the *C. pauper* group was 0.946 ± 0.011 s.e, and 3) the hybrid group was 0.50 ± 0.052 (range 0.515-0.765; Figure 1c). The number of previously assigned hybrids decreased by 47.6% in 2005 (from 21 to 11 birds) and 76.5% in 2013 (from 17 to 4 birds) using SNP data (Figure 2).

**Figure 2.**
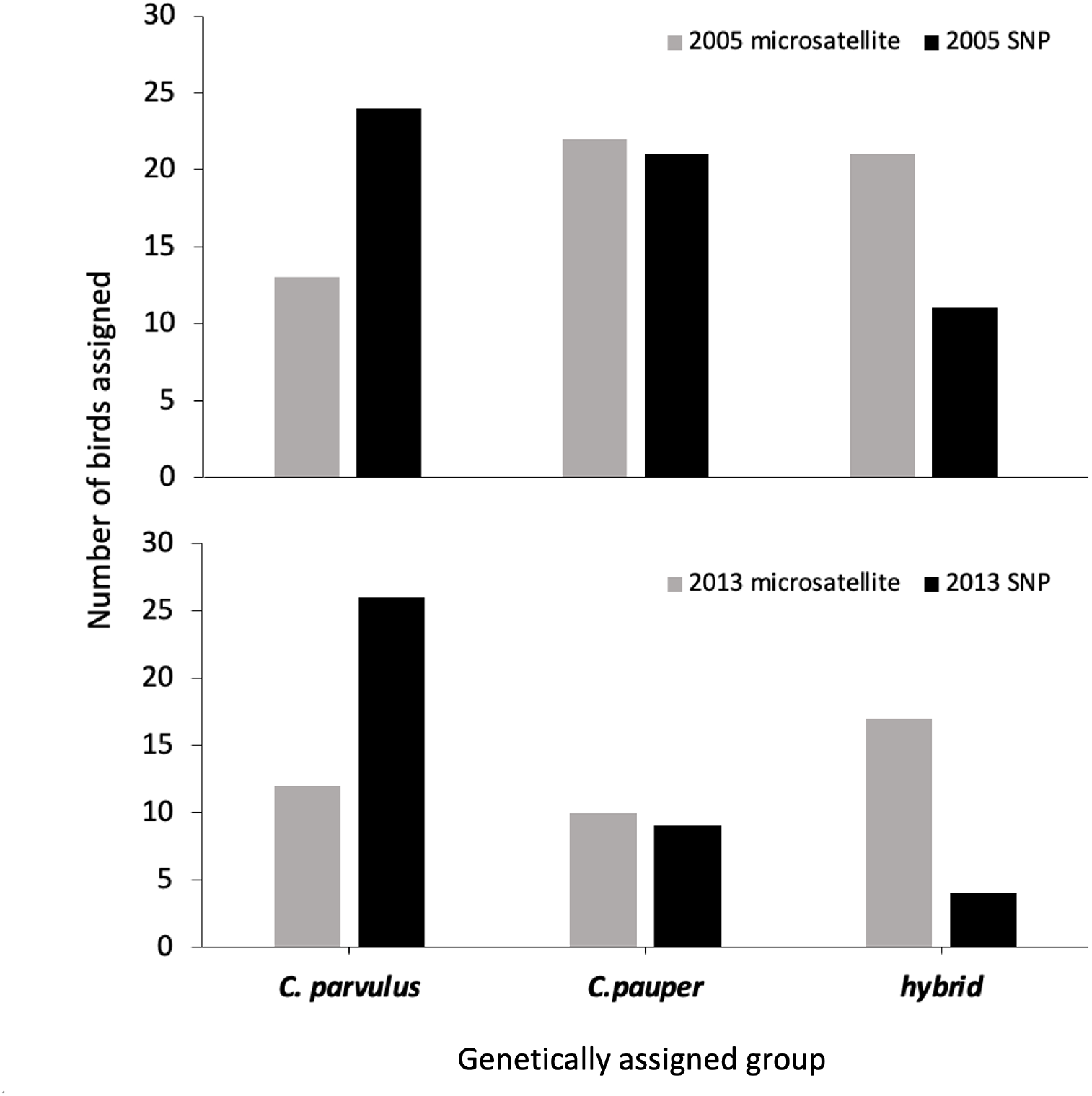
Numbers of birds assigned to each genetic group (*C. parvulus*, *C. pauper* or hybrid) using 10 microsatellites (Kleindorfer et al. 2014; Peters et al. 2017) versus 6637 SNPs used in the current study. Microsatellite assignments were based on *Structure* (Pritchard et al. 2000), SNP assignments were based on *snmf* (Frichot et al. 2014), both using a 0.80 assignment threshold value.

The results of our *fineRADStructure* coancestry matrix showed a clear separation between the two parental species groups, with *C. pauper* tending to share overall higher ancestry with one another than *C. parvulus* (Figure 2). The clustering dendrogram formed a total of 15 lineages, with 7 lineages found within the *C. parvulus* cluster and 8 within the *C. pauper* cluster.

However, some of these lineages (3 within the *C. parvulus* cluster and 2 from the *C. pauper* cluster) corresponded with hybrid individuals. Notably, we identified an instance of high co- ancestry among hybrid_3 and *C. parvulus*_21 (Figure 3) sampled in 2005. The hybrid individual had an ancestry coefficient in *snmf* of 0.7 to the *C. parvulus* group, suggesting that this individual may share recent ancestry with *C. parvulus*_21. Of note is that *C. parvulus*_21 was also an outlier with respect to having very low genetic relatedness compared to all other samples except hybrid_3 (Figure S1). In addition, individuals *C. parvulus*_15 and *C. parvulus*_12 from 2013 showed high ancestry, which is also evident in pairwise relatedness (Figure S1). Although individuals *C. parvulus_*25 and *C. parvulus*_37 also showed higher than average ancestry (Figure 2, S1), these birds were caught in 2005 and 2013 respectively, therefore recent co- ancestry is unlikely. Notably, there was a cluster of three hybrid individuals that fell outside of the *C. parvulus* or *C. pauper*-dominated clusters (Figure 3; hybrid_3 (2005), hybrid_12 (2005) and hybrid_14 (2013). Of these, hybrid_12 and hybrid_14 had almost equally shared ancestry coefficients across the two parental clusters (in *snmf*) of 0.42 and 0.52 to *C. parvulus*. All other hybrids had *snmf-*based ancestry coefficients that were between 0.69-0.77, and they were clustered within the parental genetic group to which they were more closely assigned in *fineRADstructure* (Figure 3).

**Figure 3.**
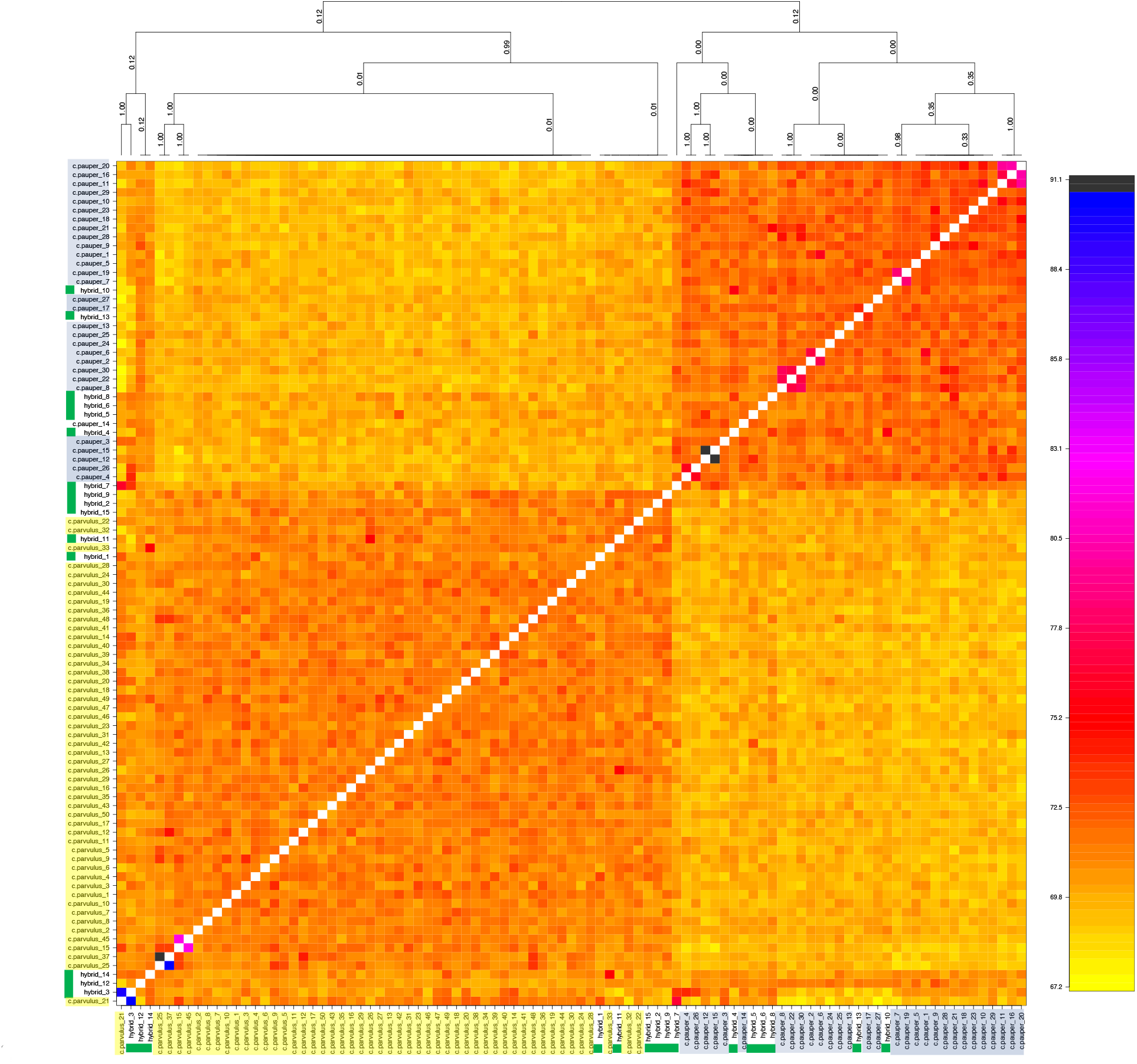
Co-ancestry matrix visualised as a heat map *fineRADstructure* and *RADpainter*. The colour of each cell in the matrix (estimated co-ancestry) shows the number of expected shared genetic markers copied from a donor genome (column) to a recipient genome (row). Support for the branches on the clustering dendrogram (co-ancestry tree) are shown on the plot. Yellow highlighted samples are *C. parvulus*, blue is *C. pauper*, and individuals with green highlighted are hybrids, as assigned by genetic clustering analysis using *snmf*.

Pairwise FST (calculated from 6757 SNPs) was highest between the two parental species (FST = 0.0353; 95% lower CI = 0.0333, upper CI = 0.0373) followed by *C. pauper* and the hybrid group (FST = 0.0095; 95% lower CI = 0.0080, upper CI = 0.0116), with a similarly low FST between *C. parvulus* and the hybrid group (FST = 0.0091; 95% lower CI = 0.0077, upper CI = 0.0116). This was also reflected in analysis of genetic relatedness between groups, with the genetic relatedness in hybrid birds being intermediate (Figure S4).

### Outlier analysis and private alleles

Both *Bayescan* and *OutFlank* FST outlier tests detected no outlier loci. However, *pcadapt* detected 69 outlier SNPs across 63 loci that had significantly divergent allele frequencies between the genetic groups identified (Figure 4, S3). Of 10463 total SNPs analysed, 4869 private alleles were found across 4167 SNPs in the parental species, with the majority being in *C. parvulus* (N = 3856; mean allele frequency = 0.09 ±0.002 s.e., min = 0.03, max = 0.52) compared to *C. pauper* (N = 1013; mean allele frequency = 0.12 ±0.001 s.e., min = 0.05, max = 0.70). These private alleles were represented across all 30 chromosomes of the *C. parvulus* genome (Figure S4-S6). A total of 348 of these parental private alleles had an allele frequency ≥0.3 within hybrid birds (mean = 0.43 ±0.006 s.e, range 0.30-1.00). Of these high frequency private alleles in hybrids, 263 (75.6%) originated from *C. parvulus*, and 85 (24.4%) originated from *C. pauper*. The majority of the 348 private alleles found in hybrids were at a higher frequency in hybrid birds than in their parental species of origin (*C. parvulus*: 253 of 263; 96.2%; *C. pauper*: 76 of 85; 89.4%). This was reduced when considering all private alleles irrespective of their hybrid frequencies, where the allele frequency in hybrids was greater than in *C. parvulus* for 72% (2773/3856) of alleles, and in *C. pauper* for 43.2% (438/1013) of private alleles.

**Figure 4.**
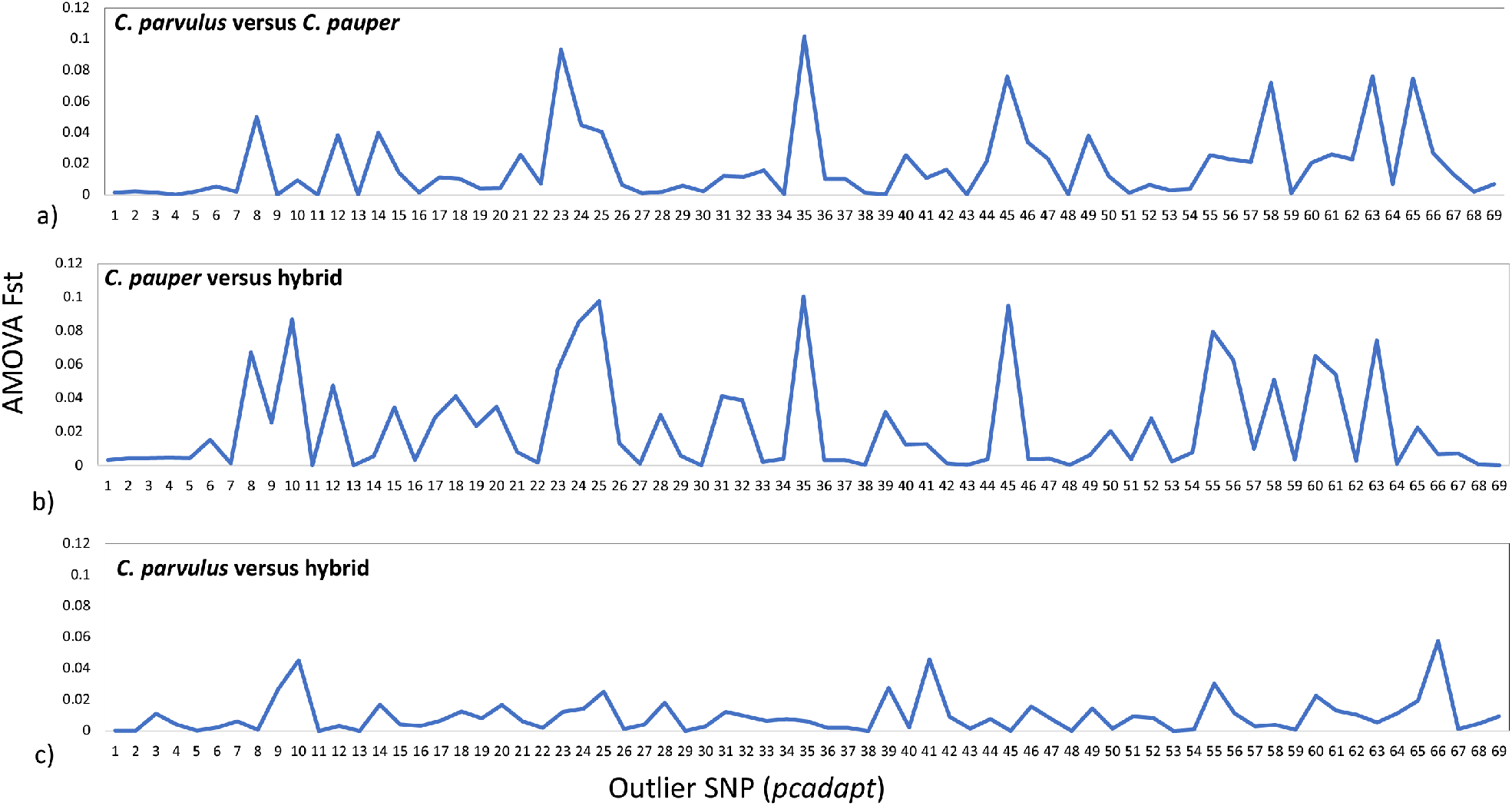
Pairwise AMOVA Fst for the 69 outlier loci identified using *pcadapt* for a) *C. parvulus* versus *C. pauper*, 2) *C. pauper* versus hybrid, and 3) *C. parvulus* versus hybrid.

### Local PCA

Genome scans using 100bp windows resulted in 417 windows analysed using a multi- dimensional scaling (MDS) approach across the *C. parvulus* genome. An aggregated PCA plot was created from MDS coordinates and two MDS axes were found to significantly explain the variation in the data (Figure S5). The three extreme window clusters identified contained a total of 63 outlier windows (Figure 5a, S6; Li and Ralph 2019). The local PCA outlier windows were mostly randomly distributed across each chromosome of the *C. parvulus* genome (Figure 5b; Li and Ralph 2019). From these 63 outlier windows, we identified 46 private alleles that were previously identified in hybrid birds with allele frequencies ≥0.3.

**Figure 5.**
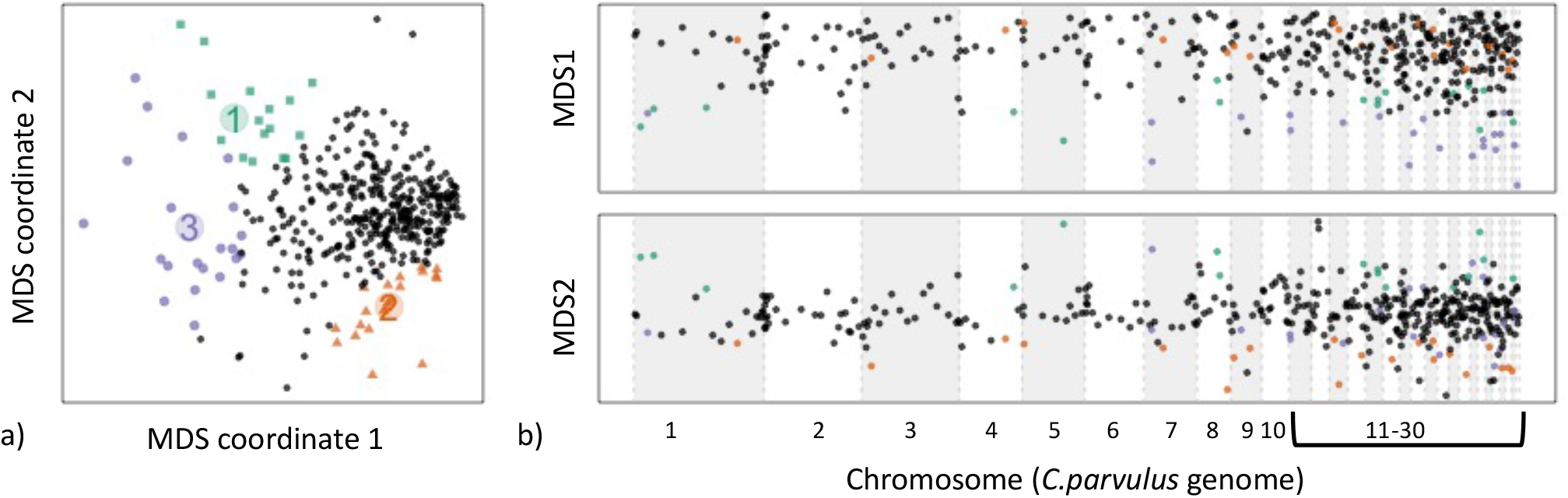
Results of multi-dimensional scaling (MDS) analysis (localPCA in *lostruct*). In a), three extreme clusters of windows are colour-coded in an aggregated PCA of MDS 1 and MDS 2 coordinates. Group 1 (green points) corresponds with *C. parvulus*, group 2 (orange) corresponds with *C. pauper, and* group 3 (purple) corresponds with the hybrid group. In b), MDS coordinates of outlier (colour-coded clusters) and non-outlier (black points) windows are shown mapped on to chromosomes of the *C. parvulus* genome (N = 30 chromosomes).

### Combined evidence for selection in hybrids across years

Between 2005 and 2013, an allele frequency increase ≥0.3 was observed in 989 private alleles (25.6% of total) originating from *C. parvulus* (mean change = 0.170 ±0.003s.e.), and 355 private alleles (35% of total) originating from *C. pauper* (mean change = 0.174 ±0.005s.e.). However, a decrease in allele frequency ≥0.3 was more commonly observed for private alleles originating from *C. parvulus* (1714 alleles, 44.5% of total; mean change = 0.128 ±0.002s.e.), which was not clearly reflected in *C. pauper* private alleles (378 alleles, 37.3% of total; mean = 0.112 ±0.005s.e.). Therefore, *C. parvulus* is contributing more private alleles to hybrids overall, but these alleles are potentially more likely to be selected against compared to *C. pauper*.

We next examined the correspondence between private allele frequency changes between 2005 and 2013, *pcadapt* outlier loci, and local PCA outlier windows. A total of 46 *pcadapt* outliers mapped to the chromosome level genome assembly for *C. parvulus*, and of these, 29 outliers aligned directly with a private parental allele found within the hybrid samples (Figure S7). These outlier private alleles each occurred on a separate chromosome so were physically unlinked and were distributed across 21 of the 30 *C. parvulus* chromosomes, with 1-6 outlier alleles per chromosome. Of these 29 private outlier alleles, 28 showed allele frequency changes from 2005- 2010, with 11 *C. parvulus* and three *C. pauper* alleles increasing in allele frequency. In contrast, 13 private outlier alleles from *C. parvulus* and one allele from *C. pauper* decreased in allele frequency between 2005 and 2013 (Figures S7). Two of the private outlier loci mapped within a local PCA outlier window on chromosomes 5 and 15 respectively, and therefore were amongst the strongest of candidates under selection, with a change in allele frequency of approximately 0.5 between years (Figures S7).

### Gene annotation

A total of 59 unique gene annotations were obtained from the 63 outlier sequences identified from *pcadapt* (Table S2). From the loci containing the 46 hybrid private alleles from the MDS outlier windows, 22 uniquely annotated genes were identified (Table S1, S3). Across all annotations, 14 loci were annotated to 2-7 variants of the same gene (Table S3). All annotations were found to be protein-coding genes or gene variants and had an average percentage match identity of 89.9% (n = 125 total annotations including variants; range: 77.9-100%). All 15 overlapping annotations were from the same locus on Chromosome 17 (RAD locus ID 695787, L761590.1). The remaining seven loci were annotated to Chromosome 4A (n=1), Chromosome 5 (n=1), Chromosome 26 (n=1) and Chromosome 28 (n=2) and the Z Chromosome (n=1).

Many gene annotations were associated with inflammation and immunity according to previous studies. For example, one of the two private outlier loci that showed an approximate 0.5 change in allele frequency between 2005 and 2010 (mentioned above) was found within an MDS outlier window could be annotated to the *C. parvulus* gene leucine rich repeat containing 40 (LRRC40), (RAD locus 18975). Within the 22 annotated outliers were other genes involved in inflammation and immunity (e.g. ankyrin repeat and SOCS box containing 12 (ASB12), E3 ubiquitin protein ligase 2 (SIAH2), pentraxin 3 (PTX3), and growth differentiation factor 15 (GDF15)), brain function (e.g., acid-sensing ion channel 2-like (LOC115914109), solute carrier family 7 member 5 (SLC7A5), DOP1 leucine zipper like protein B (DOP1B), and development (EBF transcription factor 2 (EBF2), thrombospondin type laminin G domain and EAR repeats (TSPEAR), UTP15 small subunit processome component (UTP15), as detailed further in Tables S1and S3.

## Discussion

We provide evidence that genomic introgression among Darwin’s tree finches, the Critically Endangered *C. pauper* and stable *C. parvulus*, introduces high frequency private alleles in hybrids that selection may be acting on across years. Alleles that were unique to each parental species appeared in hybrid birds at frequencies higher than within their parental species. Many of these alleles showed a substantial change in allele frequency over an 8-year period, while some were identified as candidate outliers under selection, or appeared in divergent genomic windows (Figure S7). Some of these alleles were located within genes that code for functions that may be relevant for hybrid fitness. Therefore, the retention of genetic diversity via hybridisation in this population of *Camarhynchus* may be beneficial for the adaptive capacity of *C. pauper*, which is under increasing threat from local extinction. Our findings further elucidate the limitations of microsatellite-based hybrid assignment in closely related and sympatric species with greater certainty afforded by high resolution, genome-wide SNP markers. Darwin’s finches offer an unparalleled opportunity to study the earliest stages of species hybridisation in a vertebrate (Grant and Grant 2008, 2014), and here they reveal the role of introgression for maintaining genetic variation of endangered species within populations.

### Mismatches in hybrid identification

Our results suggest that the number of hybrid tree finches on Floreana are overestimated using microsatellite markers (Kleindorfer et al. 2014, Peters et al. 2017). We document a much lower hybrid assignment rate using SNPs yet confirm a swarm of backcrossed *C. parvulus* and hybrid finches, which presents a challenge for classifying individuals based on genetic clustering approaches. Notably, our results do not provide further insight into the shift in the frequency of hybridisation across years that was previously documented in this population (Kleindorfer et al. 2014) because we analysed a non-random sample of birds that was pre-selected based on prior microsatellite assignments. Furthermore, this study does not negate the findings of female-biased introgression from *C. pauper* to *C. parvulus* in Peters et al. (2017), as their conclusions were based on genetic assignment probabilities to each parental species rather than distinct groups of hybrids and parental individuals. We found that the number of hybrid birds assigned using SNP data was 47.6% lower in our 2005 sample (from 21 to 11 birds) and 76.5% lower in our 2013 sample (from 17 to 4 birds) compared to the initial microsatellite-based assignments (Peters et al. 2017). Notably the co-ancestry approach of *fineRADstructure* grouped just three hybrid birds outside of the two parental clusters, and these all had ∼0.5 assignment probability to each parent, indicating potential first-generation hybrids without backcrossing (Figure 3). The rest of the hybrid-assigned birds had *snmf* assignment probabilities to each parental cluster that were lower or higher than 0.5. The detection of birds with hybrid ancestry in *fineRADStructure* therefore appears to be conservative, perhaps influenced by the recent ancestry of the parental species. Of note is that in the current study we use a different genetic clustering approach implemented in the R package *snmf,* while previous studies used the program *Structure* (Pritchard et al. 2000), though we use the same assignment cut-off value of 0.80 in this study as in Peters et al (2017).

This difference in program usage is not likely to have resulted in the large differences we report in hybrid assignments between the SNP and microsatellite analyses, particularly as we used three approaches to determine hybrid status (e.g., PCA, *snmf*, and *fineRADStructure*).

For genetic structure analyses, previous studies of threatened species that have compared SNP and microsatellite data have found similar patterns, but SNP data often provide stronger evidence of hierarchical genetic structure, and increased evidence for population evolutionary independence (Zimmerman et al. 2020), with genetic assignments sometimes varying across lineages (Bohling et al. 2019). Marker comparisons for detecting hybrid animals have found microsatellites to overestimate hybridization frequencies compared with high resolution SNP markers that have a much higher density and wider coverage of the genome (i.e., in newts: Miralles et al. 2024; polecats: Szatmári et al. 2021). SNPs may also reveal higher resolution of backcrossed individuals that were missed by microsatellites (e.g., among deer and sika: MacFarlane et al. 2019; birds: Väli et al. 2010). Overall, this suggests that SNP approaches outperform microsatellite-based assessments of hybridization frequency.

Our analyses support the ‘hybrid swarm’ scenario observed in Peters et al. (2017), whereby a continuum of hybrid ancestry exists between the two parental tree finch species, though we find a reduced proportion of hybrids in our dataset than previously reported (Figure 2, Kleindorfer et al. 2014). This suggests that there are fewer tree finch hybrids in the Floreana population than previously thought, which may be promising for maintaining the genetic integrity of endangered *C. pauper*. Notably, hybrid recruitment in the tree finch population on Floreana increases in years with higher rainfall (Kleindorfer and Dudaniec 2020), thus additional sampling periods would enable the impact of climate fluctuations to be accounted for in estimates of hybridisation frequency. Although we advance present understanding using a RADseq approach, our study could be further improved with a whole genome sequencing approach that enables a higher density genome scan of diversity and selection, as well as structural variants that may be under selection in hybrids.

### Private alleles and selection in hybrids

Theory predicts that genome-wide patterns of introgression in hybridising populations will vary and is influenced by either population size variation (Currat et al. 2008) or selection (Martin and Jiggins 2017). We find widespread evidence for introgression across the genome (using the *C. parvulus* reference genome), with private alleles in hybrids, outlier loci and divergent genome windows located on nearly all 30 *C. parvulus* chromosomes, rather than being clustered within specific genomic regions. Therefore, it is unlikely that we have missed hotspots of introgression across the finch genome. Most of the 348 high frequency (≥0.30) private alleles found in hybrids had a lower frequency in their parental species of origin, indicating that either genetic drift or selection in hybrids is driving this difference. With 70.6% of private hybrid alleles being retained and undergoing substantial allele frequency change over the eight-year sampling interval, a case for selection is strengthened by the outlier loci and outlier windows that overlap with the private alleles, as identified using *pcadapt* and local PCA, respectively (Figure S7).

Given the cryptic hybridization found in this group of birds, as in many other species groups, this study will aid future work in identifying hybrids based on the presence of private alleles. Further, through the identification of private alleles, this study will aid in genetically identifying ‘pure’ members of the endangered *C. pauper* on Floreana and help to understand positive or negative roles that hybridisation may play for this tree finch population. Indeed, hybridisation is increasingly heralded as a conservation tool rather than a hindrance (Chan et al. 2019).

Accordingly, we previously found that nests of hybrid *Camarhynchus* birds on Floreana harbour fewer parasites of the avian vampire fly, *Philornis downsi*, compared to nests of parental species (Peters et al., 2019), suggesting that hybridization confers some resistance to *P. downsi*, which in turn may favour greater genetic introgression due to heterosis (Peters et al., 2019; Peters & Kleindorfer, 2015; Peters & Kleindorfer, 2018). The increasing numbers and co-evolutionary shifts of *P. downsi* in Floreana finch nests have been well documented over the past two decades (Kleindorfer and Dudaniec 2016; Common et al. 2019, 2023), alongside rates of tree finch hybridisation (Kleindorfer et al. 2014; Peters et al. 2019). Further, hybrid birds are reported to be much more common on Floreana Island than on the island of Daphne Major, where *P. downsi* is in low numbers, and hybrids occur in just 2-5% of breeding pairs (Grant & Grant, 2008; 2016).

As introgression should favour alleles beneficial for hybrid fitness, the ancestry of beneficial genotypes within hybrids is expected to show an excess of ancestry towards the introgressing species (Fraïsse et al. 2022; Martin and Jiggins 2017). In Darwin’s tree finches on Floreana Island, the more common and stable *C. parvulus* is contributing more of its genetic material to hybrid birds than the critically endangered *C. pauper* via backcrossing from hybrids to *C. parvulus (*Peters et al. 2017). Our findings support this, with *C. parvulu*s private alleles being more common in hybrids (Figure S7). These *C. parvulus* private alleles were also more likely to show substantial allele frequency change across the eight-year sampling period (24 *C. parvulus* versus 4 *C. pauper* alleles; Figures S7). Thus, the pattern of introgression we observe indicates variation in retention or selection for private alleles originating from *C. parvulus,* while alleles from *C. pauper* remain more stable. Several striking examples demonstrate adaptive introgression during invasions of novel species into a location, or for congeneric species in sympatry or allopatry. For example, Valencia-Montoya et al. (2020) discovered the transfer of pesticide resistance alleles to invasive moths from native moths via genomic introgression.

However, introgression can have positive, negative or neutral effects and evolutionary forces acting within hybrid populations may effectively "filter" gene flow between species (Martinsen et al. 2001). Introgressed DNA is typically thought to be deleterious in the recipient species (Veller et al. 2023), therefore investigation into the incidence of purging of deleterious alleles over our study period may further elucidate the genetic mechanisms underlying this hybridisation.

### Gene annotations to outlier loci and private alleles

When selection acts on hybridising species, genes linked to reproductive isolation tend to resist introgression, while alleles that confer locally adaptive traits do not (Martin and Jiggins 2017). Here we found candidate loci under selection containing private alleles in hybrid tree finches, and that also mapped to genes associated with inflammation and immunity, brain function, and development (Table S3). The gene with the highest query cover (100%) and percentage match identity (100%) was DTX4, which is involved in notch signalling (DTX4) and is part of the pathway network for cytosolic sensing of pathogen associated DNA, and the innate immune system (Wang et al. 2021). Notably, DTX4 was found to be differentially expressed in urban and non-urban common kestrels (*Falco tinnunculus*; Damiani et al. 2024). Other genes of note that had lower query cover and percentage identification included E3 ubiquitin protein ligase 2 (SIAH2), which is known to suppress growth of avian reovirus, an important virus of chickens (Chen et al. 2018). Pentraxin 3 (PTX3) is involved in the anti-inflammatory immune response and may be used as an acute phase protein marker to monitor inflammatory conditions in poultry (Burkhardt et al. 2019). Ninjurin 1 has well-known nerve regenerative and anti-inflammatory effects, but is not studied in birds (Liu et al. 2021). For genes involved in cognition, the acid sensing ion channel (LOC115914109) is involved in detecting extracellular acidification in the brain and aids in both chemosensation and mechanosensation for monitoring homeostasis and pathological signals (Cheng et al. 2018). The gene for solute carrier family 7 member 5 (SLC7A5) has been widely studied in chickens in the context of nutrient transport, metabolism and digestion (Fagundas et al. 2020), but the gene has also been associated with feather pigmentation and follicle development in ducks (Sun et al. 2023). In development, the gene EBF2 transcription factor is involved in ligament formation and limb development in chickens (Mella et al. 2004). These gene annotations to private alleles in hybrids that are also candidate loci under selection suggest that introgression among *Camarhynchus* confers functional genetic effects that may be further investigated for fitness correlates.

## Conclusion

Our study provides evidence that natural hybridization can provide important genetic variation that impacts both ecological and evolutionary processes. We document frequent introgression in Darwin’s tree finches on Floreana Island that is promoting gene flow and novel selection pathways in hybrids that may be influencing the evolutionary trajectory of *Camarhynchus* on the Galapagos Islands. Notably, the island of Floreana is undergoing a massive ecological restoration effort (described in Kleindorfer et al. 2021) involving avian predator removal and species re- introductions, which may shift selection pressures and hybridisation frequencies for *Camarynchus* going forward. Our study therefore has implications for monitoring the genetic diversity and integrity of the Critically Endangered medium tree finch (*C. pauper*) as Floreana transitions into a state of ecological restoration.

## Supporting information

Supplementary Information_Dudaniec et al.

## Acknowledgements

On title page

## Data Archiving Statement

Data for this study are available at: to be completed after manuscript is accepted for publication.

---

**Table 1.**
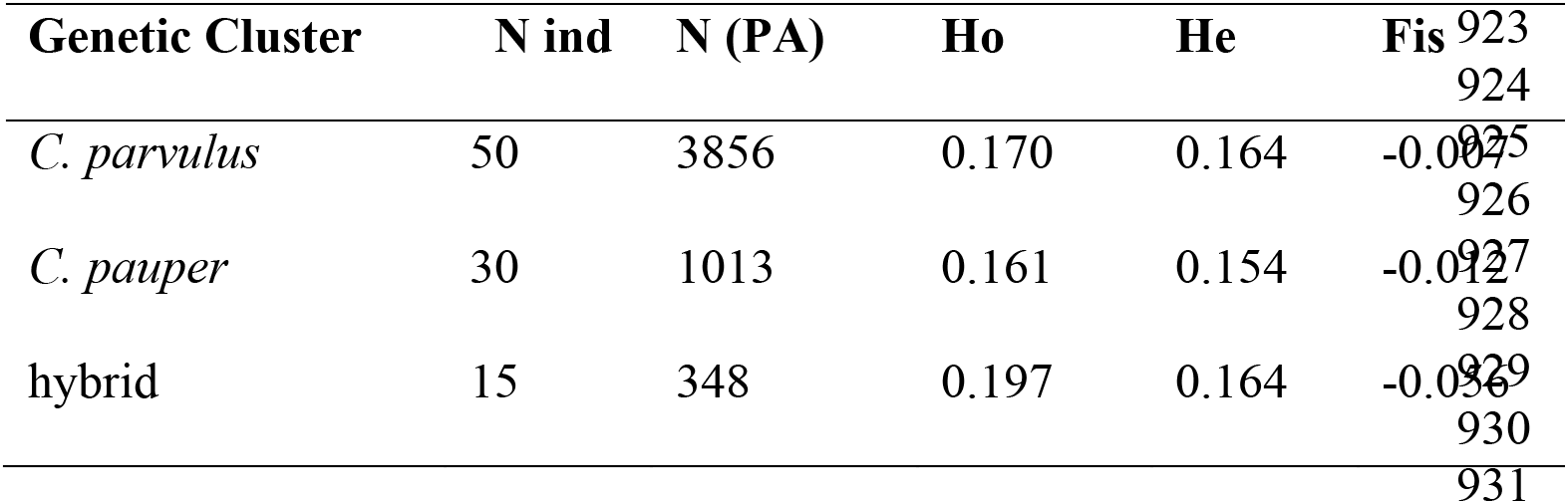
Summary statistics for parental and hybrid Darwin’s tree finches following SNP-based genetic assignment, including number of individuals (N ind), number of private alleles (N (PA)), observed heterozygosity (Ho), expected heterozygosity (He), inbreeding coefficient (Fis). Note that private alleles in the hybrid group are those that were found in both *C. parvulus* or *C. pauper*.

## Notes

### Competing Interest Statement

The authors have declared no competing interest.

