## Supplementary Information_Dudaniec et al. for "Genomic introgression between critically endangered and stable species of Darwin’s tree finches on the Galapagos Islands"

**Text S1. RAD library preparation and sequencing**

Genomic DNA was digested with the restriction endonuclease *SbfI* and processed into RAD a library similar to the method of Baird, et al., 2008. Genomic DNA was digested for 60 min at 37 °C in a 15 μL reaction with 20 units (U) of *Sbf I* (New England Biolabs, Inc., Ipswich, Massachusetts, USA). After digestion, samples were heat-inactivated for 20 min at 80 °C followed by addition of 2.0 μL of 1 nM P1 Adapter(s), a modified Solexa adapter (Illumina, Inc.). P1 adapters each contained a unique multiplex sequence index (barcode), which is read during the first ten nucleotides of the Illumina sequence read. 1 uM P1 adaptors were added to each sample along with 10x T4 DNA Ligase Buffer (Enzymatics, Inc), high concentration T4 DNA Ligase (Enzymatics, Inc), and 0.8 μL H_2_O which was then incubated at room temperature for 20 min.

Samples were heat-inactivated again at 65 °C for 20 minutes, pooled, and randomly sheared with a Bioruptor (Diagenode) to an average size of 500 bp. Samples were then run out on a 1.5% agarose, 0.5X TBE gel, and DNA 250 bp to 800 bp was isolated using a MinElute Gel Extraction Kit (Qiagen). End Repair / dA-Tailing module (NEB) was used to polish the ends of the DNA. After subsequent purification, 1 μL of 1 μM P2 adapter, a divergent modified Solexa adapter (Illumina, Inc.), was ligated to the obtained DNA fragments at RT. Samples were again purified and eluted in 15 μL. The eluate was quantified using a Qubit Fluorometer with the dsDNA HS assay kit (Invitrogen) and ~200 ng of this product was used in 40 50 μL PCR amplifications with 25 μL 2x Phusion Master Mix (NEB), 2.5 μL of 10 μM modified Solexa Amplification primer mix (Illumina, Inc.) and up to 22.5 μL H_2_O. 200 μL of the amplification product was cleaned and run on a 1.5% agarose (Sigma), 0.5X TBE gel and DNA 350 bp to 900 bp was excised and purified as before. The library was quantified with a Qubit fluorometer and run on an Agilent Bioanalyzer with the High Sensitivity kit to determine size distribution. 150 bp paired-end sequencing was performed on the Illumina HiSeq 4000.

**Text S2. RAD data processing and SNP characterisation**

A single RAD library (N = 95 samples) was demultiplexed and cleaned (-q), rescuing barcodes (-r) using the process_radtags component of *Stacks v2.54* (Rochette et al. 2019, Catchen et al. 2011, 2013, 2018). Only high-quality reads containing unambiguous barcodes and RAD cut site remnants were retained, resulting in 305.8 million (M) reads total, with a range of 1.3M – 23M reads (6.6M mean, ±412,000 SE) per sample (Table S1). Read counts varied due to differences in DNA or library quality but all samples were included in the analysis.

Paired-end reads were then analysed *de novo* by the denovo_map.pl *Stacks* pipeline. The analysis was conducted twice, before and after the samples were reassigned to their parental species or hybrid groups (see main text for details). For both assemblies, denovo_map.pl was executed allowing up to six mismatches between single-end loci (-M 6 -n 6) following optimization. The gstacks component of the denovo_map.pl pipeline was set to have a strict genotyping p-value (--gt-alpha 0.01) and PCR duplicates were removed (--rm-pcr-duplicates). Due to DNA quality, all samples had a large number of PCR duplicates (mean value of 84%); after filtering the coverage was reduced to a per-sample scaled mean of 4.6x (1.9 – 12.4x) resulting in the assembly of 69k loci on average per sample (28K – 103K) (Table S1). The assembly cumulatively produced 379,399 RAD loci with a mean length of 300.3 bp and a consistent phasing was found for 93.9% of those loci (indicating high fidelity). Across the metapopulation 369,530 single-nucleotide polymorphisms (SNPs) were identified from the RAD loci. Further filtering of SNP markers was done within the *populations* program in *Stacks* as described in the main text.

For analyses that required a reference genome, the consensus sequences from the loci in the *de novo* catalog (catalog.fa.gz) were aligned against the *Camarhynchus parvulus* reference genome STF-HiC (1.28 GB; Genbank accession number: GCF_901933205.1) using BWA mem (Li 2013). The stacks-integrate-alignments program was then used to inject the alignment coordinates back into the *de novo* assembled Stacks output files, allowing for reference-based analyses to be executed.

*Private allele analysis*

To conduct the private allele analysis, using the reference-aligned data, the populations program was run including only individuals from the two parental species to identify private alleles within either species. This was undertaken after analysis of genetic structure and genetically assigning hybrids and parental species. The populations program was executed again, this time including only the genetically-assigned hybrids and exporting a VCF file defining the SNPs found within the hybrids. The summary statistics file (which flags private alleles), along with the hybrid VCF export, were provided to the stacks-private-alleles program which plotted parental private alleles, found in the hybrids, along each *C. parvulus* chromosome, according to its frequency in the hybrid birds. These values were smoothed using a kernel-smoothing algorithm included in the stacks-private-alleles program.

**Table S1 –** Excel file “Table S1 RADseq Sample Statistics.xls’ - Statistics of RAD sequencing filtering and number of SNPs.

**Table S2 –** Excel file “TableS2_pcadapt69Annotations” - Full annotation list of the 69 pcadapt outlier loci with results from BLASTN mapped to *C. parvulus* reference genome.

**Table S3.** Gene annotations (NCBI BLAST) for loci with private alleles in hybrid birds that were found within MDS outlier windows against the *Camarhynchus parvulus* reference genome using BLASTN. ‘Perc. Ident’ = percentage match identity. Annotations with a ‘*’ are variants of the same gene. Loci with ‘†’ were identified as a candidate outlier under selection using *pcadapt*.

| **Locus** | **Chr** | **Annotation** | **Query cover** | **E value** | **Per. Ident** | **Accession ID** |
| --- | --- | --- | --- | --- | --- | --- |
| 572730 | 5 | deltex E3 ubiquitin ligase 4 (DTX4) | 75% | 0 | 100 | XM_030949919.1 |
| 655258 | 4A | family with sequence similarity 155 member B (FAM155B) | 100% | 0 | 100 | XM_030967768.1 |
| 724872 | 26 | complement component receptor 1-like protein (LOC115913499) | 30% | 3.00E-74 | 100 | XM_030965544.1 |
| 728068 | 28 | growth differentiation factor 15 (GDF15) | 15% | 7.00E-36 | 98.82 | XM_030966952.1 |
| 728011 | 28 | kelch like family member 26 (KLHL26), transcript variant X2 | 17% | 3.00E-40 | 97 | XM_030966698.1 |
| †695787 | 17 | DOP1 leucine zipper like protein B (DOP1B) | 9% | 6.00E-17 | 96.36 | XM_030949668.1 |
| †695787 | 17 | pentraxin 3 (PTX3), mRNA | 9% | 7.00E-16 | 96.23 | XM_030954812.1 |
| †695787 | 17 | *ATPase phospholipid transporting 9A (putative) (ATP9A), transcript variant X2 | 21% | 1.00E-33 | 89.08 | XM_030963283.1 |
| †695787 | 17 | *ATPase phospholipid transporting 9A (putative) (ATP9A), transcript variant X1 | 21% | 1.00E-33 | 89.08 | XM_030963282.1 |
| †695787 | 17 | ninjurin 1 (NINJ1) | 23% | 5.00E-37 | 89.06 | XM_030956942.1 |
| †695787 | 17 | ankyrin repeat and SOCS box containing 12 (ASB12) | 24% | 7.00E-36 | 87.68 | XM_030967669.1 |
| 695787 | 17 | E3 ubiquitin protein ligase 2 (SIAH2) | 22% | 7.00E-31 | 86.51 | XM_030954336.1 |
| †695787 | 17 | SH3GL interacting endocytic adaptor 1 (SGIP1) | 14% | 3.00E-15 | 85.37 | XM_030953810.1 |
| †695787 | 17 | acid-sensing ion channel 2-like (LOC115914109) | 24% | 7.00E-31 | 85.29 | XM_030966463.1 |
| †695787 | 17 | solute carrier family 7 member 5 (SLC7A5) | 24% | 6.00E-27 | 84.21 | XM_030955834.1 |
| †695787 | 17 | acrosin-like (LOC115910452) | 19% | 2.00E-21 | 84.07 | XM_030960625.1 |
| †695787 [ | 17 | pterin-4 alpha-carbinolamine dehydratase 1 (PCBD1) | 25% | 2.00E-26 | 82.86 | XM_030951320.1 |
| †695787 | 17 | G protein-coupled receptor 21 (GPR21) | 20% | 4.00E-18 | 82.3 | XM_030961208.1 |
| †695787 | 17 | *uncharacterized LOC115913567 (LOC115913567), transcript variant X2, ncRNA | 25% | 4.00E-23 | 81.43 | XR_004061391.1 |
| †695787 | 17 | *uncharacterized LOC115913567 (LOC115913567), transcript variant X1, ncRNA | 25% | 4.00E-23 | 81.43 | XR_004061390.1 |
| †695787 | 17 | *TBC1 domain family member 16 (TBC1D16), transcript variant X3 | 32% | 2.00E-31 | 81.22 | XM_030962295.1 |
| †695787 | 17 | *TBC1 domain family member 16 (TBC1D16), transcript variant X2 | 32% | 2.00E-31 | 81.22 | XM_030962294.1 |
| †695787 | 17 | *TBC1 domain family member 16 (TBC1D16), transcript variant X1 | 32% | 2.00E-31 | 81.22 | XM_030962293.1 |
| †695787 | 17 | thrombospondin type laminin G domain and EAR repeats (TSPEAR), transcript variant X1 | 33% | 4.00E-33 | 81.18 | XM_030954492.1 |
| †695787 | 17 | EBF transcription factor 2 (EBF2), transcript variant X2 | 15% | 4.00E-08 | 80.23 | XM_030964056.1 |
| 536713 | Z | *UTP15 small subunit processome component (UTP15), transcript variant X2 | 37% | 2e--30 | 79.5 | XM_030969100.1 |
| 536713 | Z | *UTP15 small subunit processome component (UTP15), transcript variant X1 | 37% | 2e--31 | 79.5 | XM_030969099.1 |

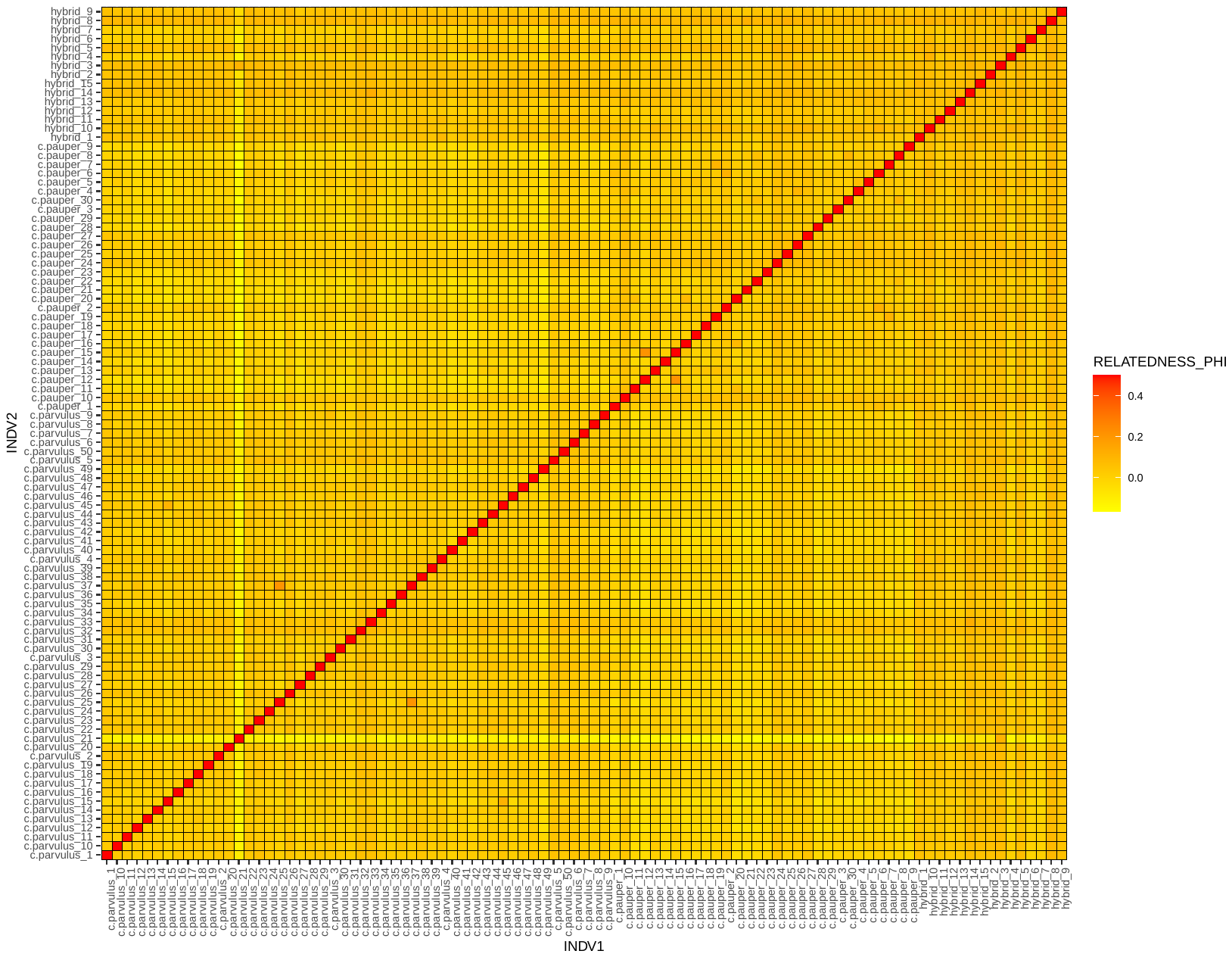

**Figure S1.** Pairwise genetic relatedness heatmap between all tree finch samples (N = 95, sampled 2005 and 2013) using relatedness2 in R. *C. parvulus_21 (*individual 964*)*, was identified as having very low relatedness to all other samples except for hybrid_3. Higher than average relatedness was found for individual *C. parvulus*_25 with *C. parvulus*_37, and also for *C. pauper*_15 with *C. pauper*_12, and *C. parvulus*_15 with *C. parvulus*_45.

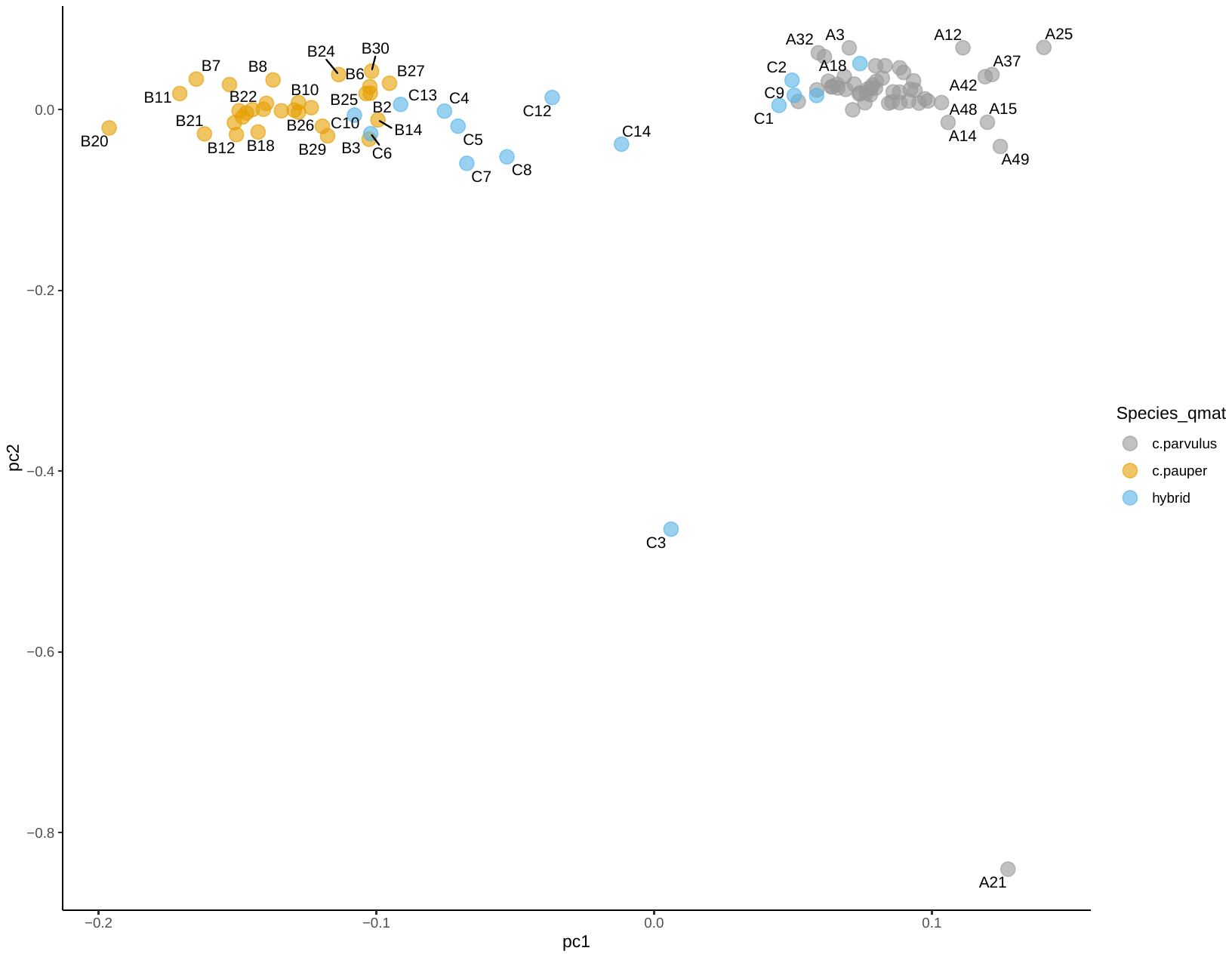

A)

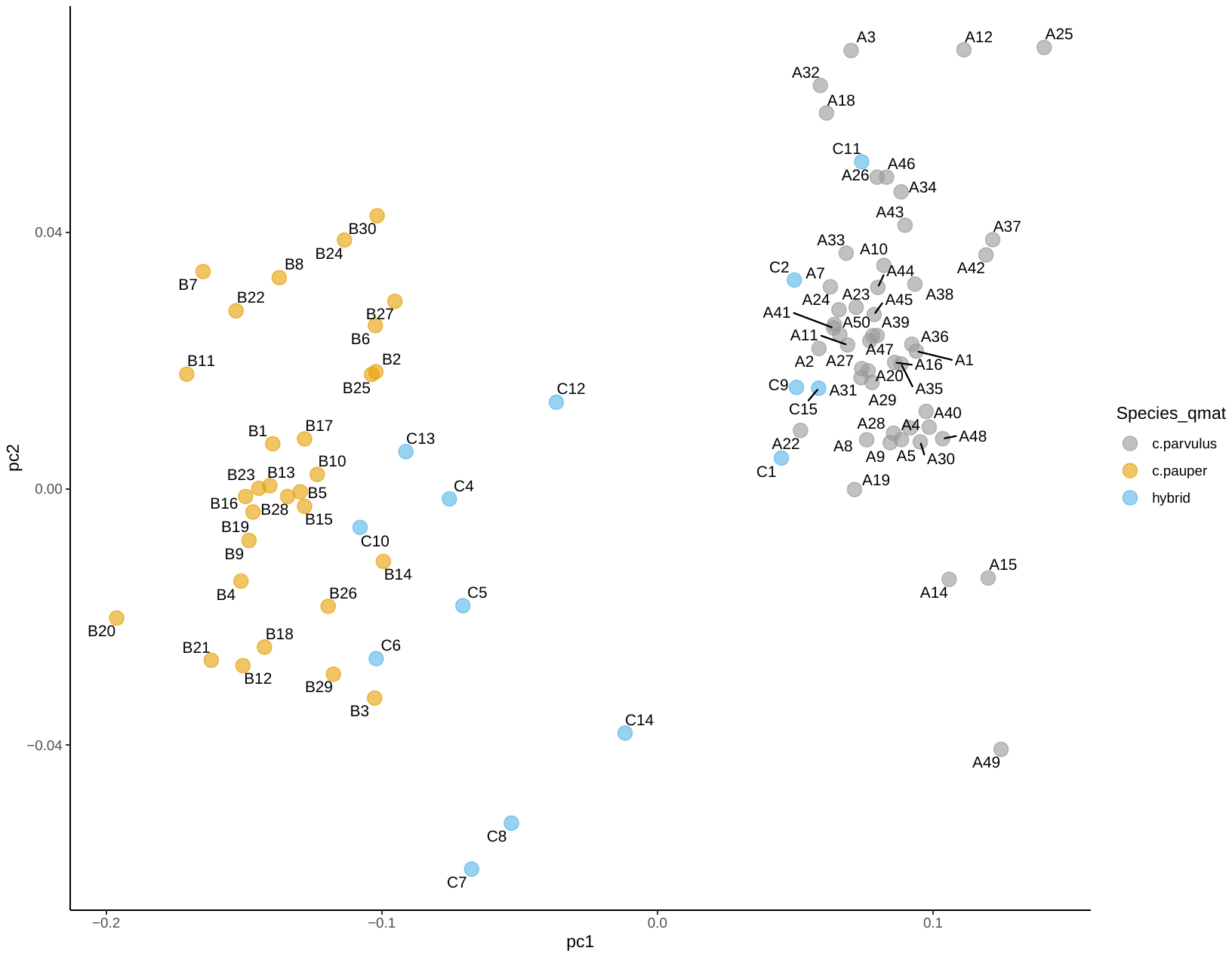

B)

**Figure S2.** PCA plot showing samples labelled according to SNP-based assignments to parental (*C. parvulus* and *C. pauper* in yellow and grey) and hybrid (in blue). A) shows PCA plot (PC1 and PC2 axes) for all samples (N = 95), B) shows the PCA plot excluding the outliers C3 (hybrid_3) and A21 (*C. parvulus*_21), which showed high co-ancestry in fineRADstructure. *C. parvulus*_21 further showed very low pairwise relatedness to all other samples except for hybrid_3 (see Figure S2).

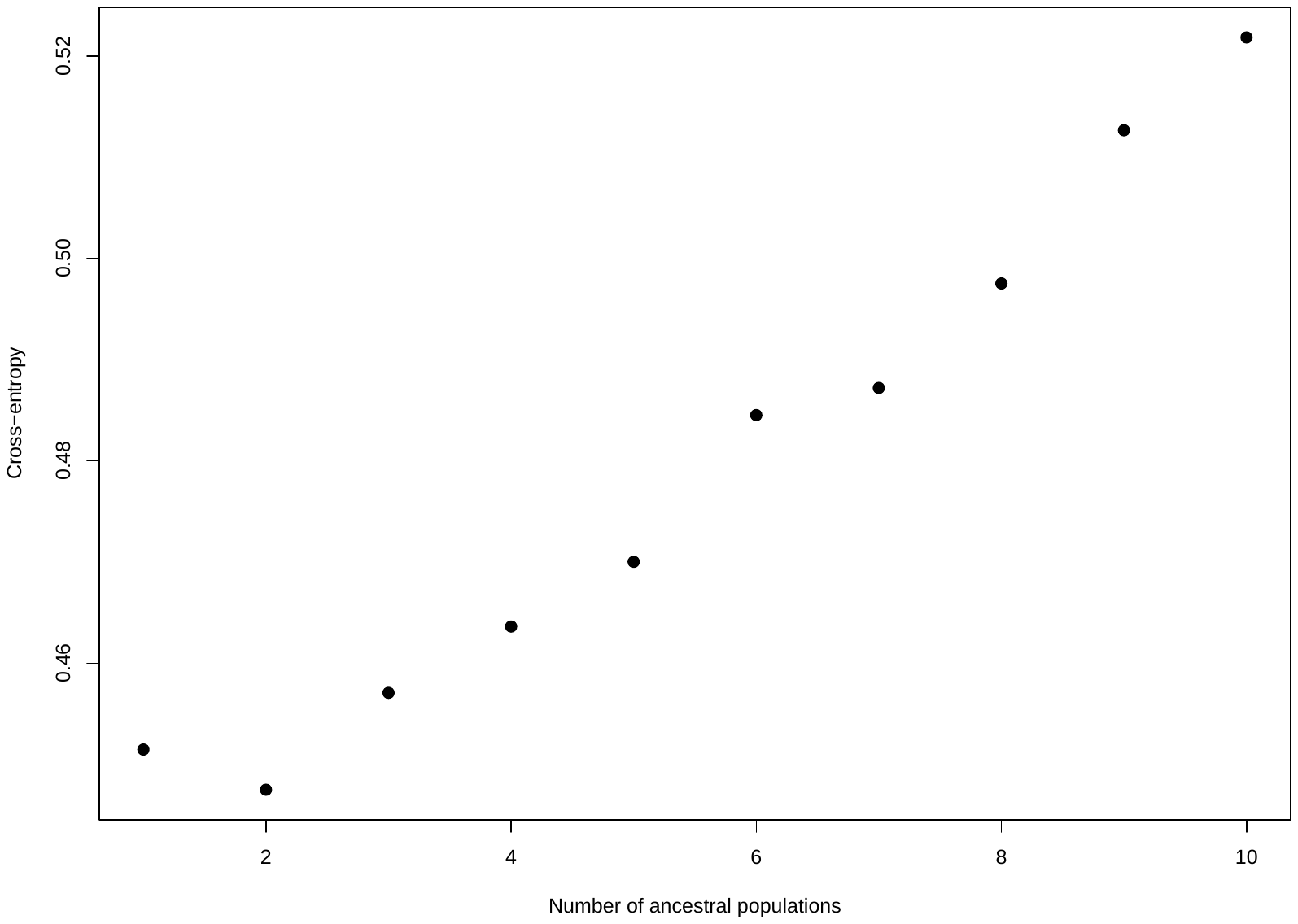

Figure S3: Snmf cross-entropy values showing most supported number of clusters is 2.

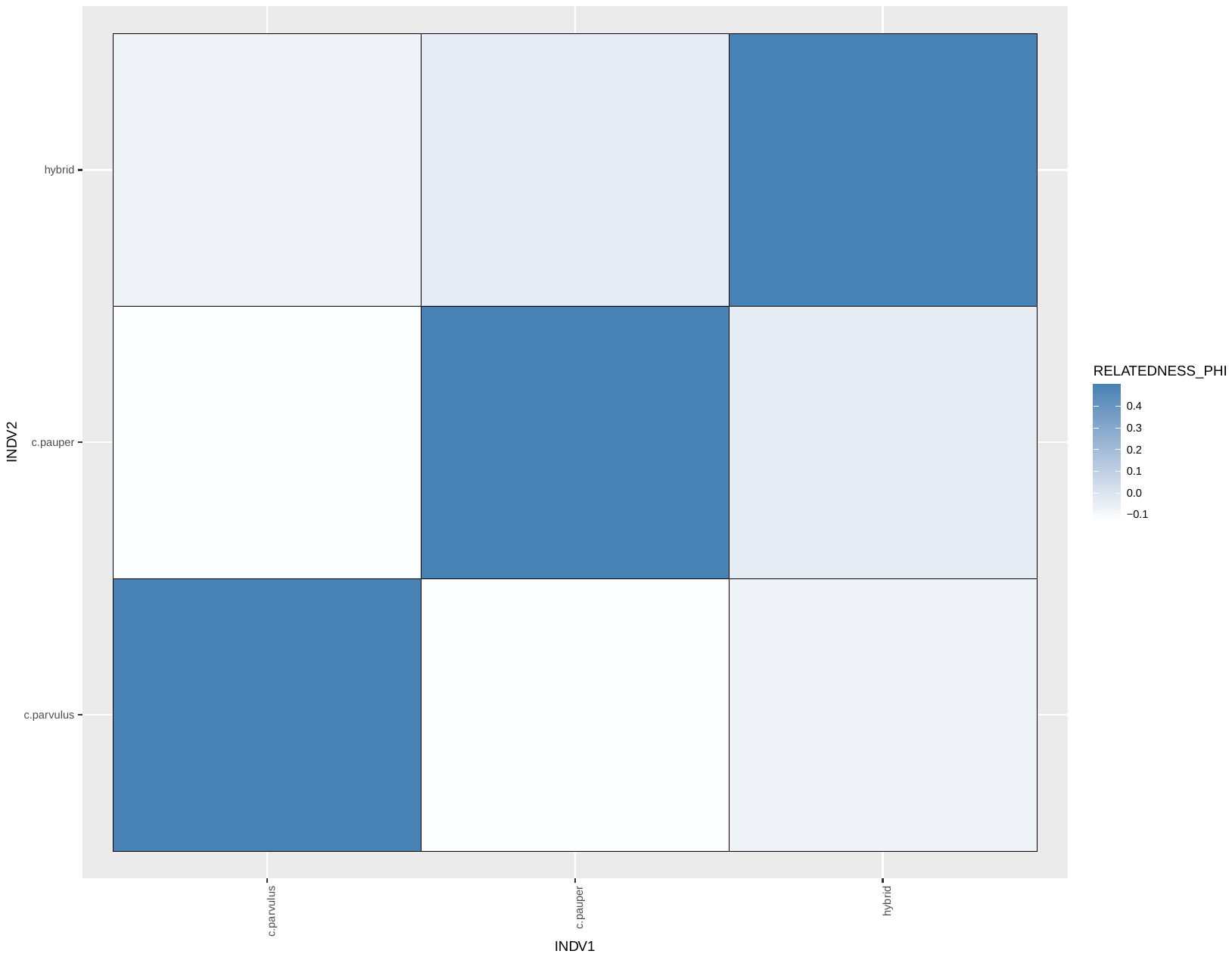

**Figure S4.** Pairwise genetic relatedness between the SNP-assigned parental species and hybrid groups.

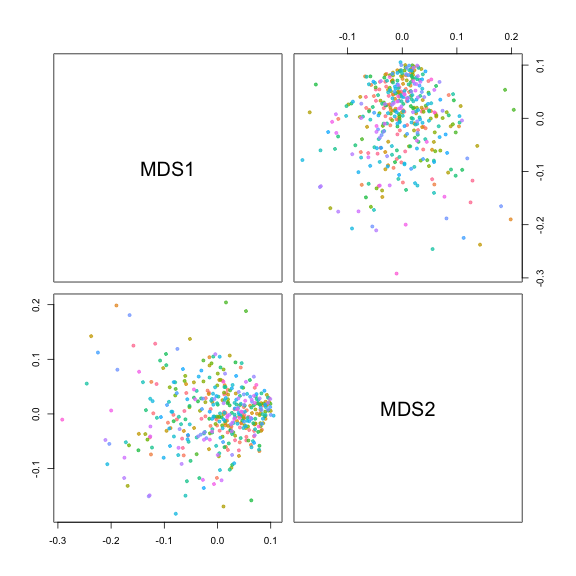

**Figure S5.** The two multi-dimensional scaling (MDS) axes found to significantly explain variation in the data, coloured by chromosome.

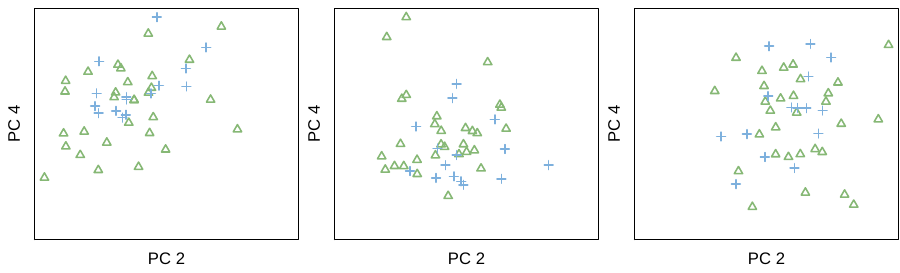

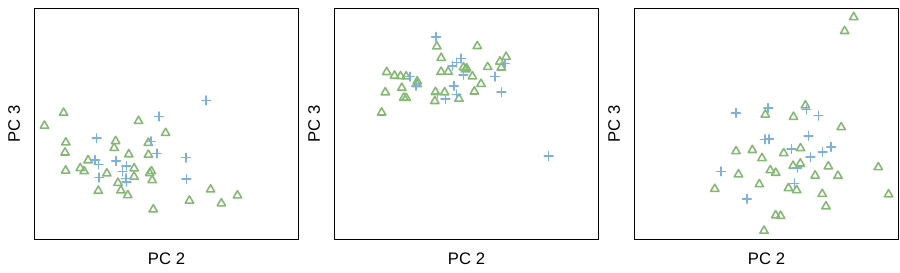

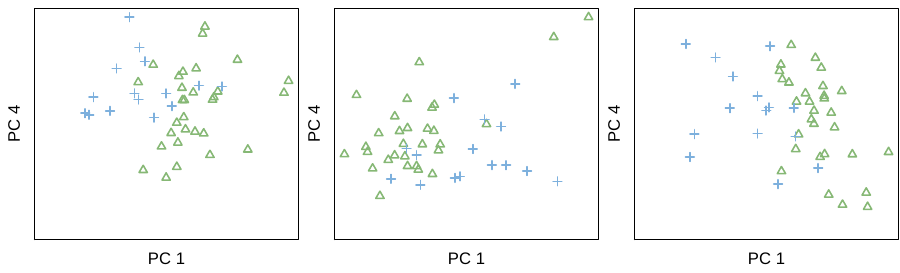

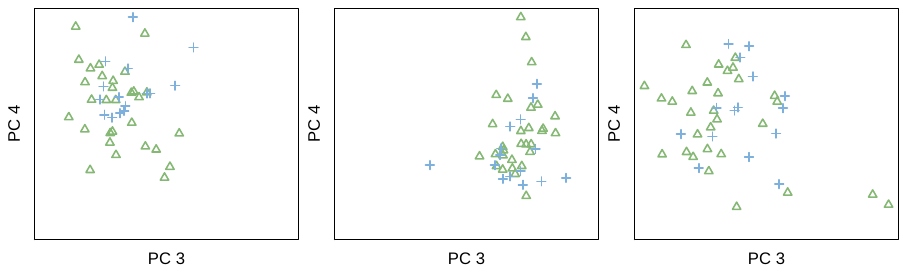

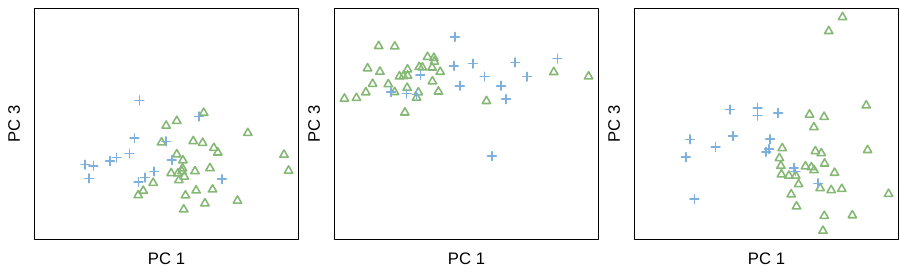

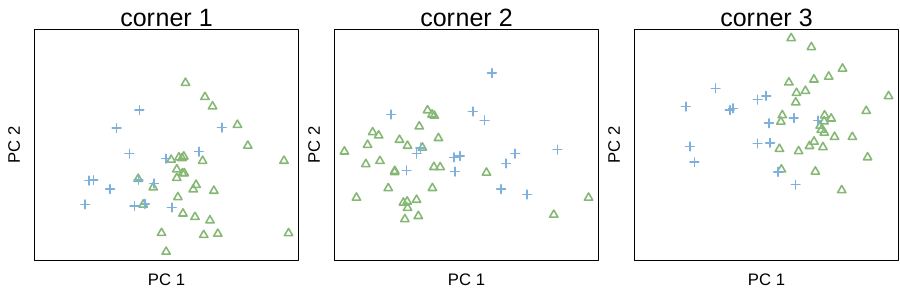

**Figure S6.** Principal components axes 1-4 combinations for the three corners identified using multi-dimensional scaling (MDS), coloured by the three identified window clusters.

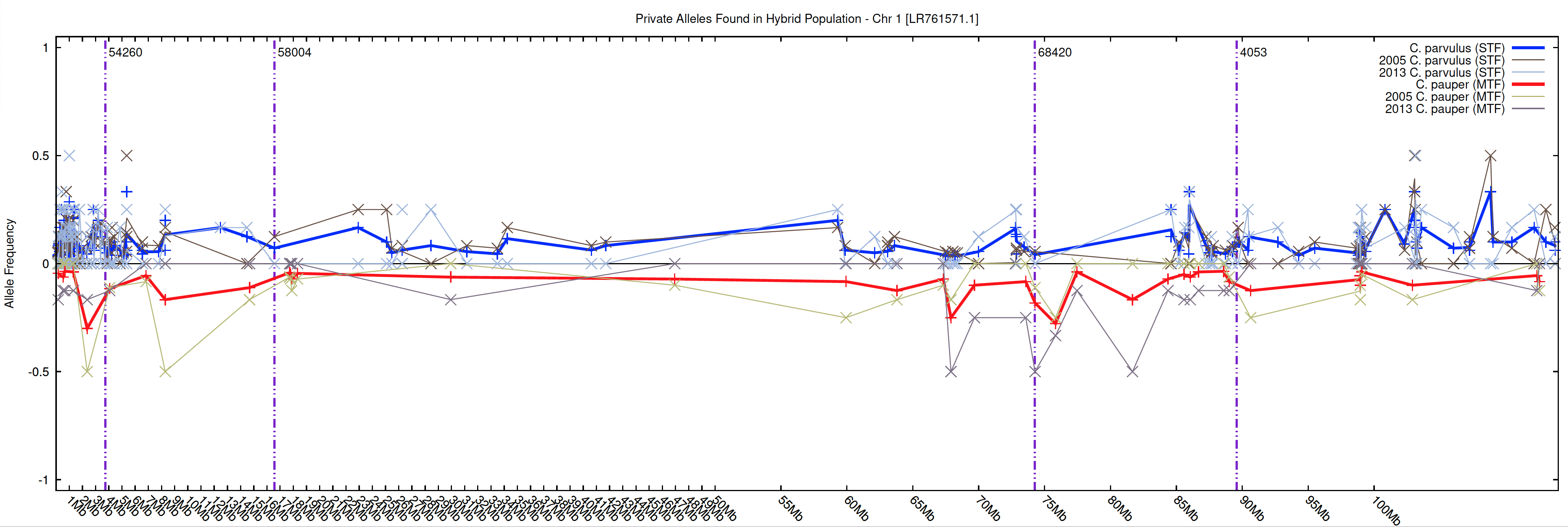

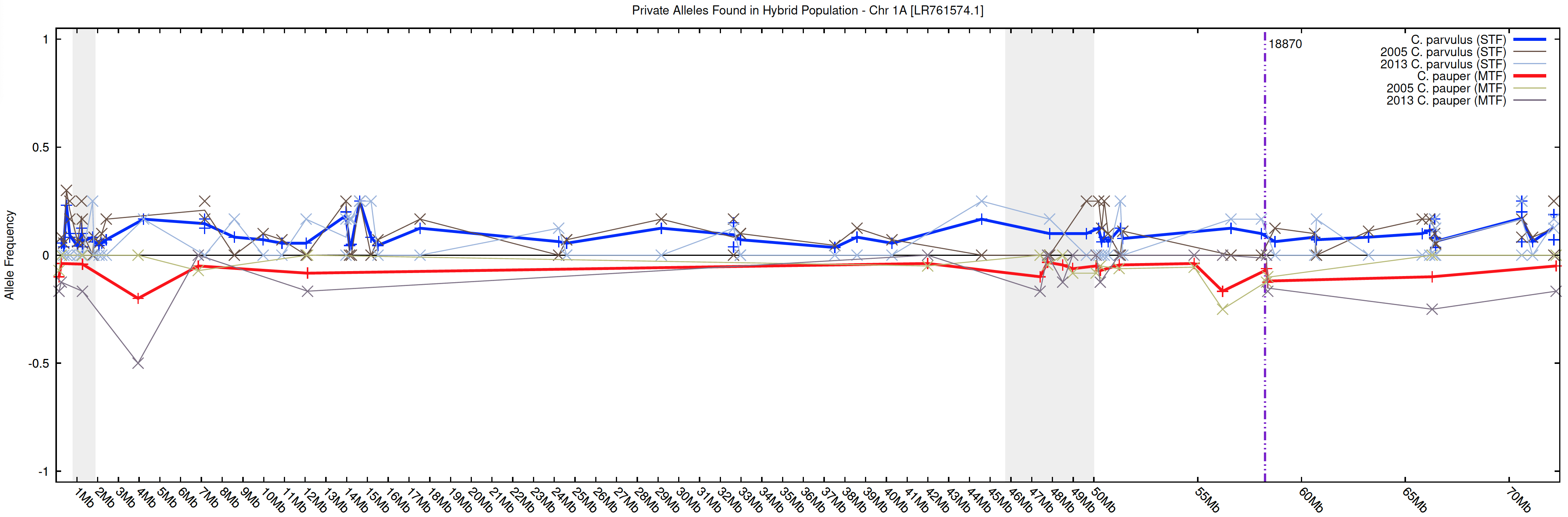

2

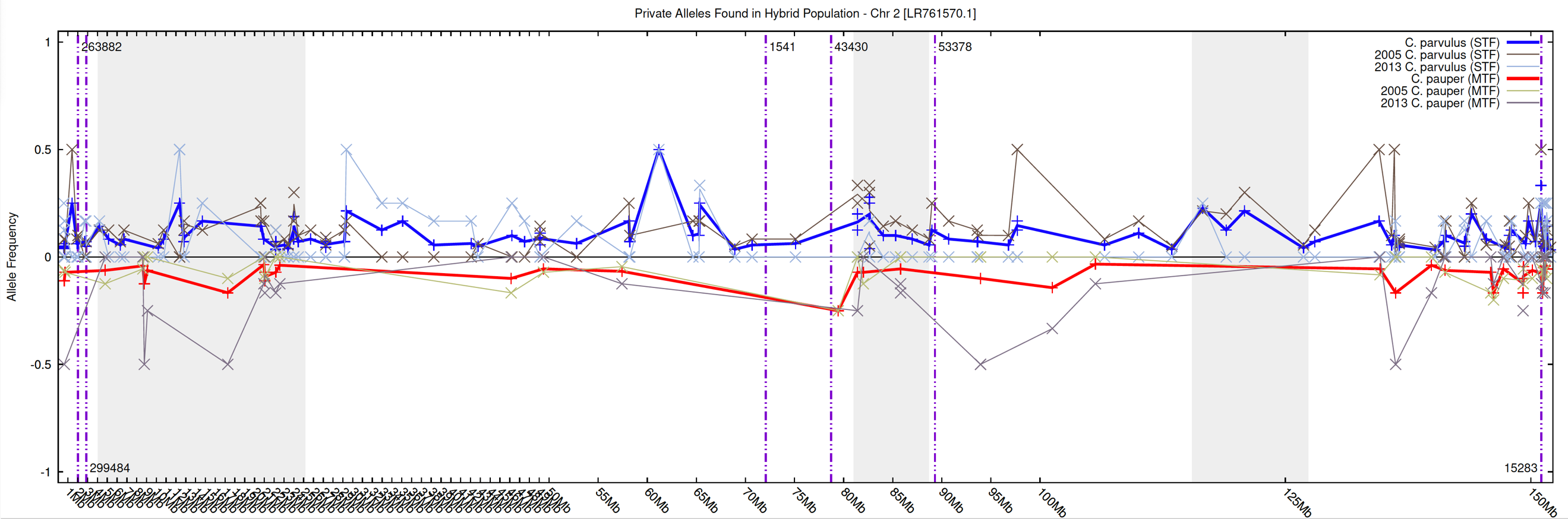

3

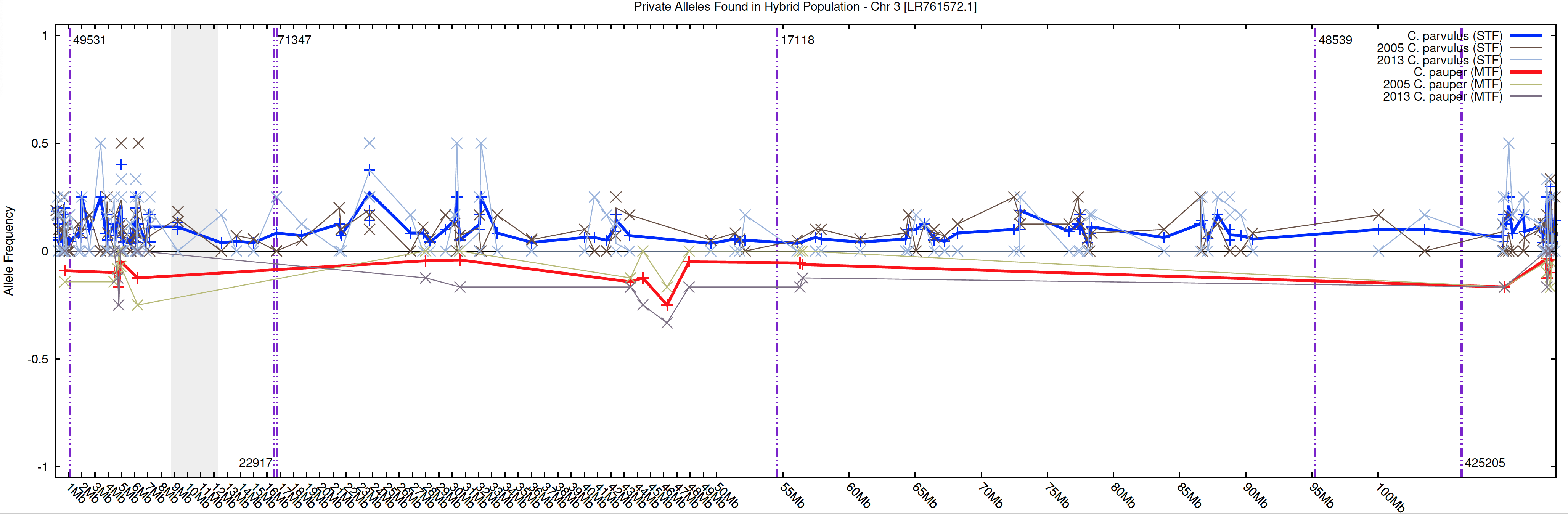

4

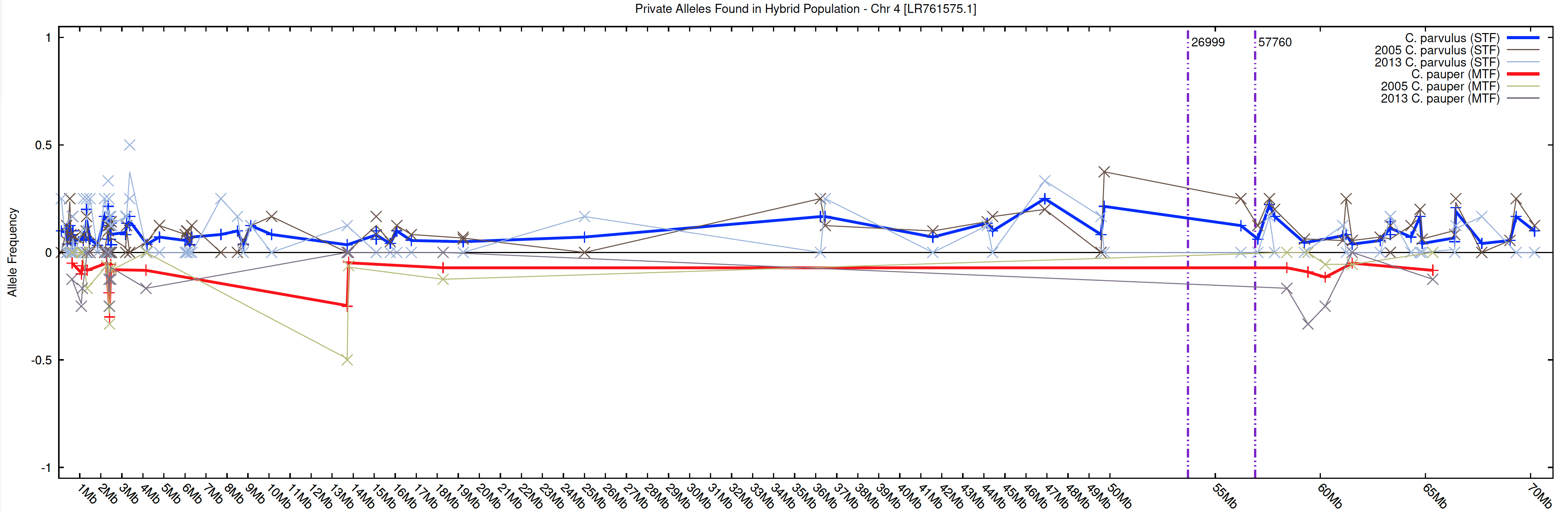

5

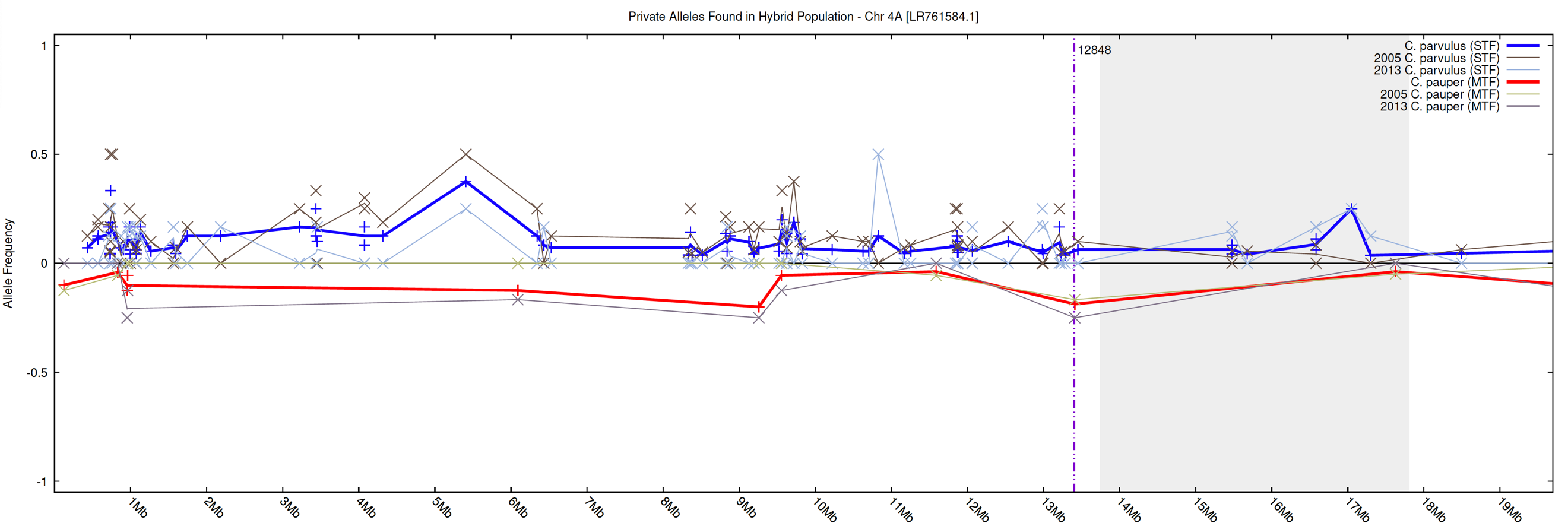

6

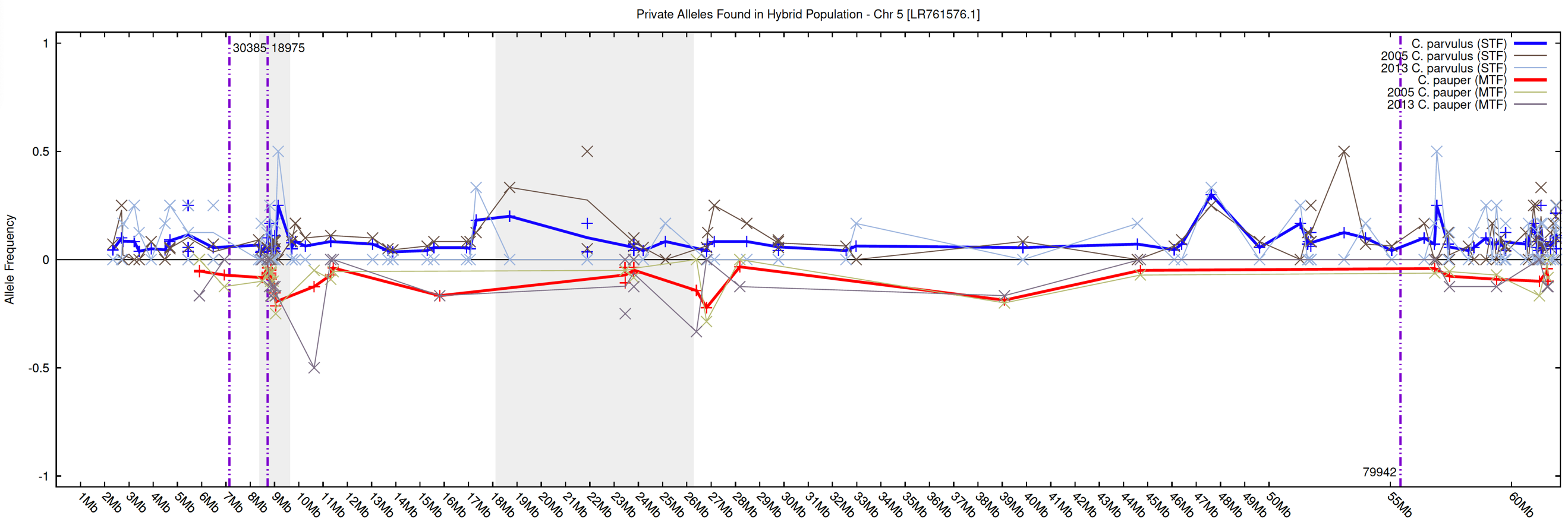

7

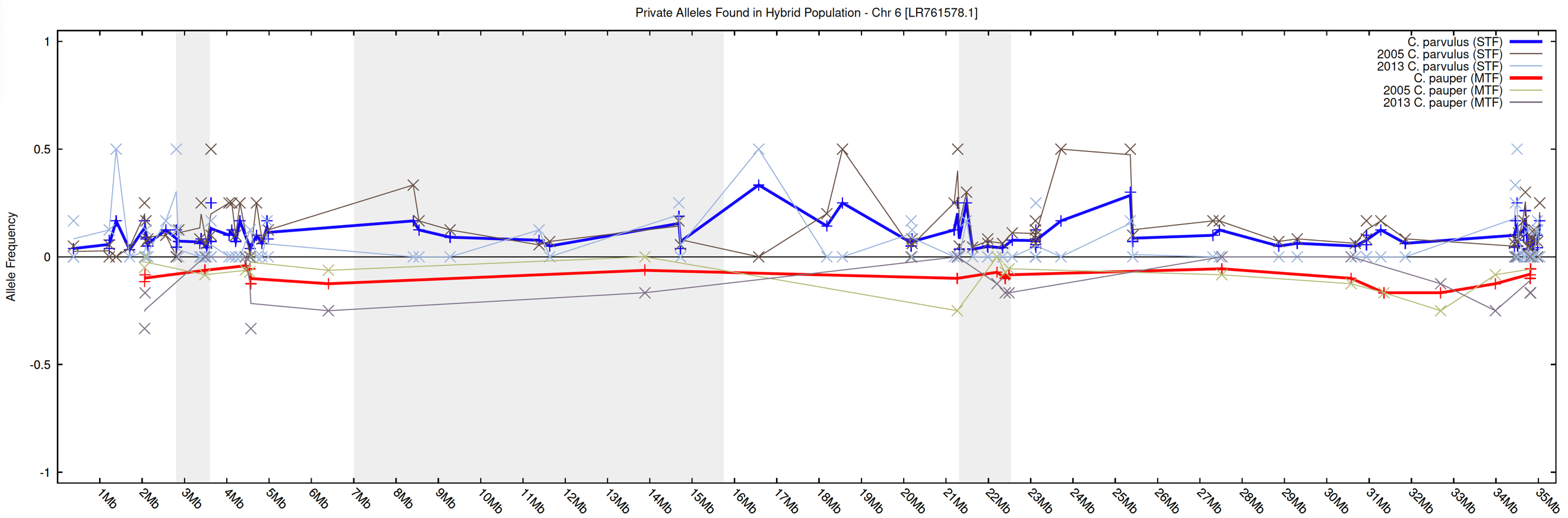

8

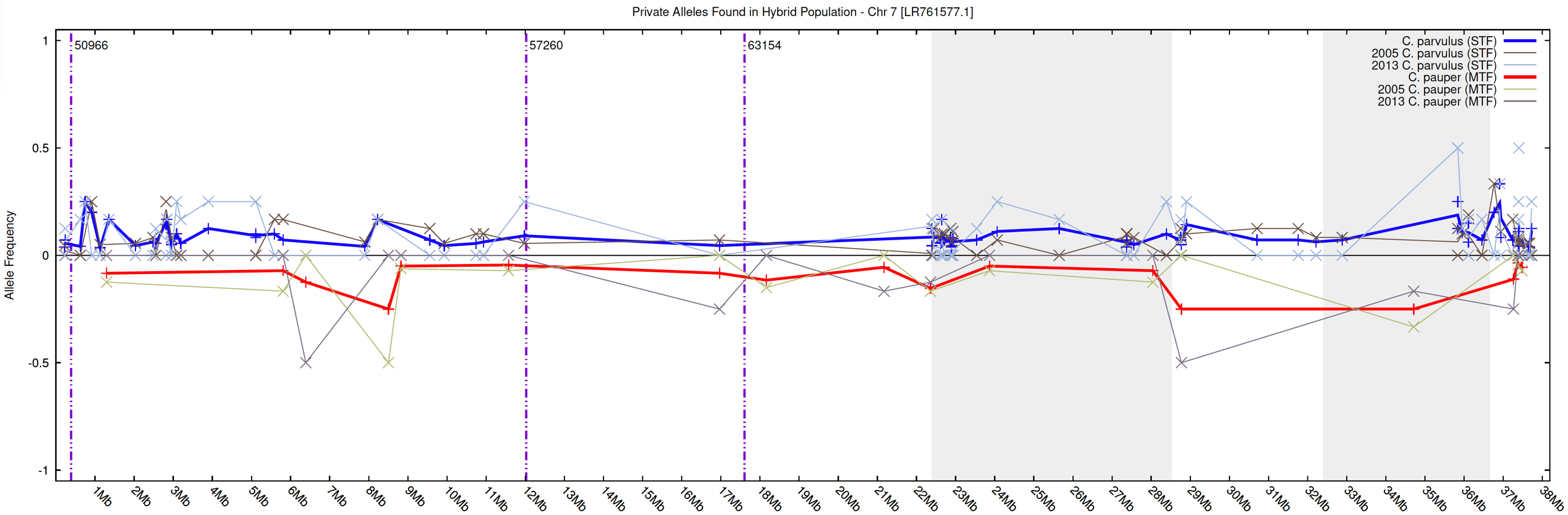

9

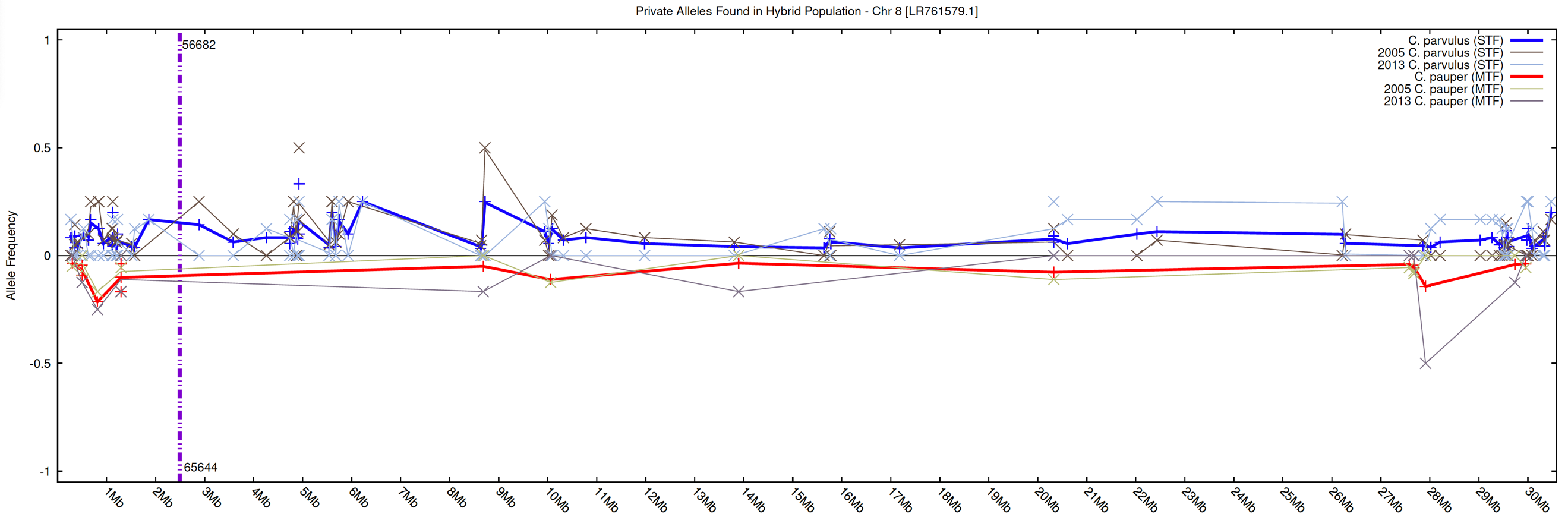

10

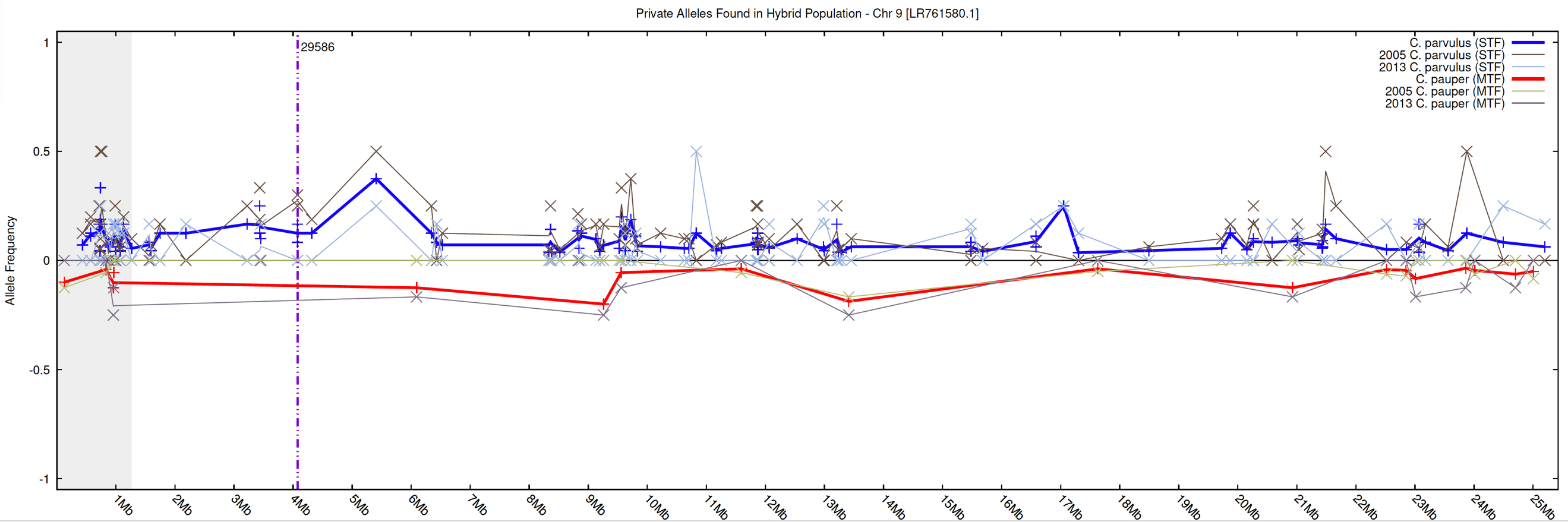

11

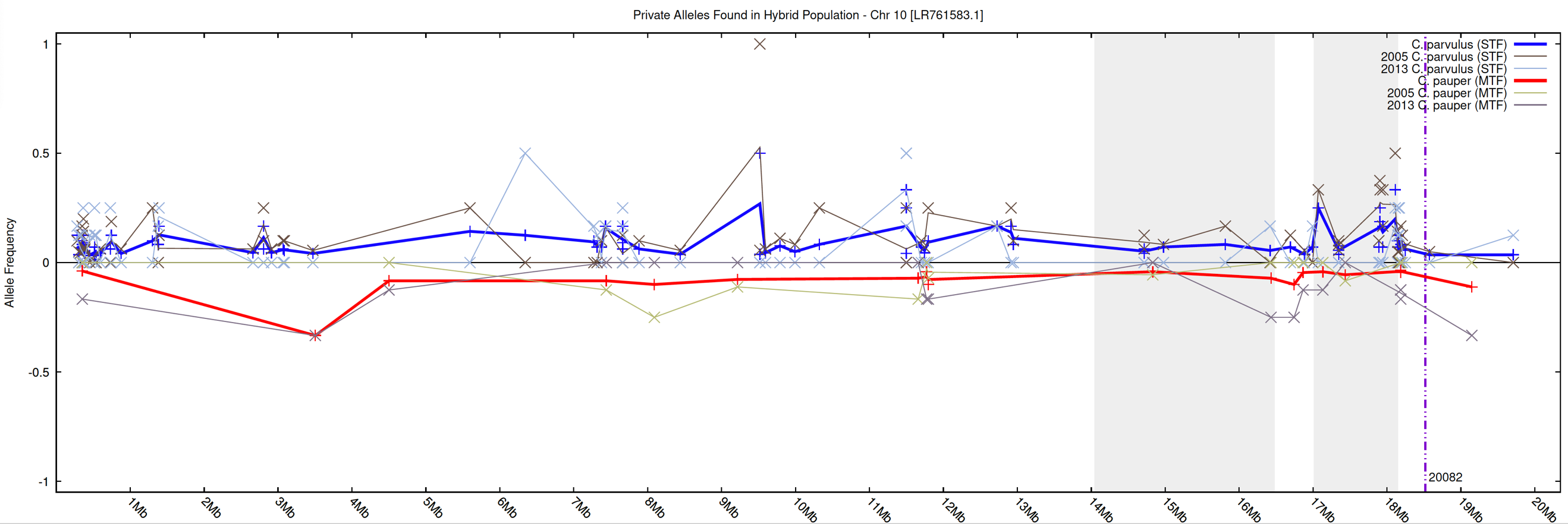

12

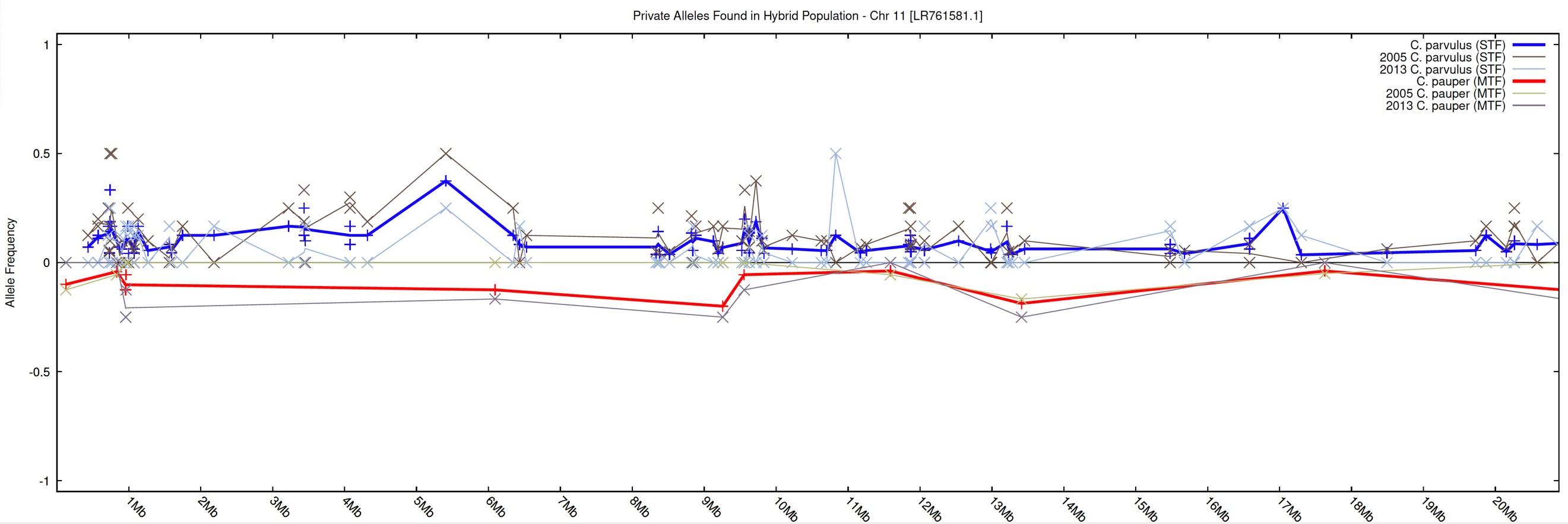

13

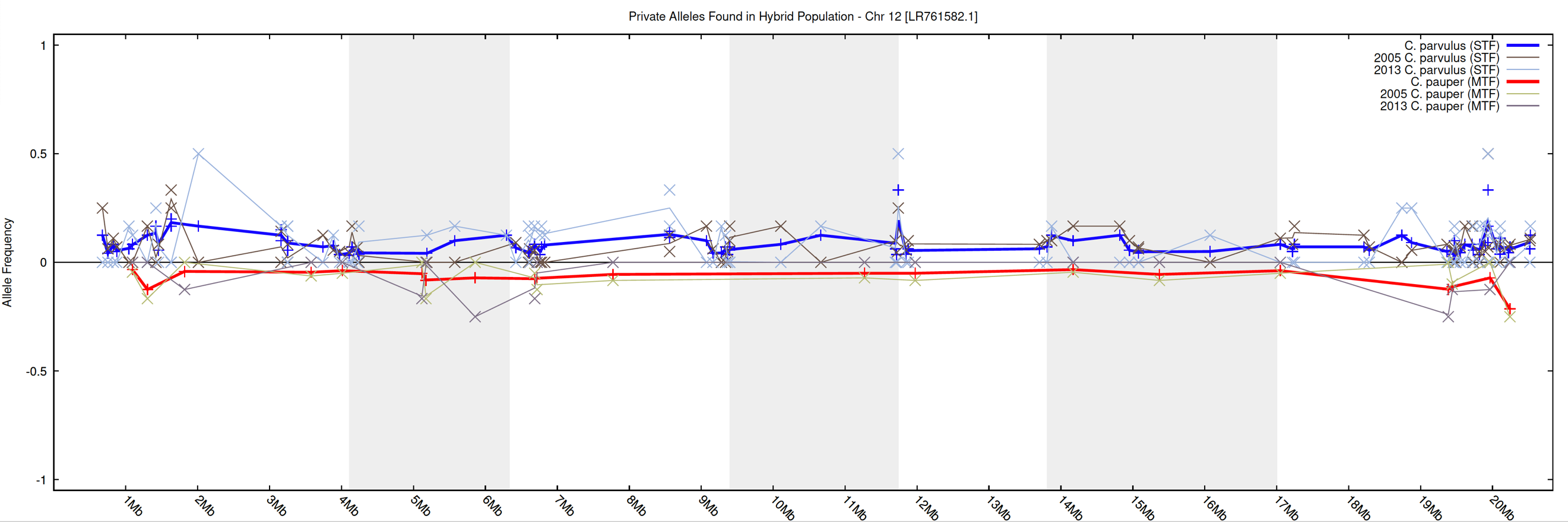

14

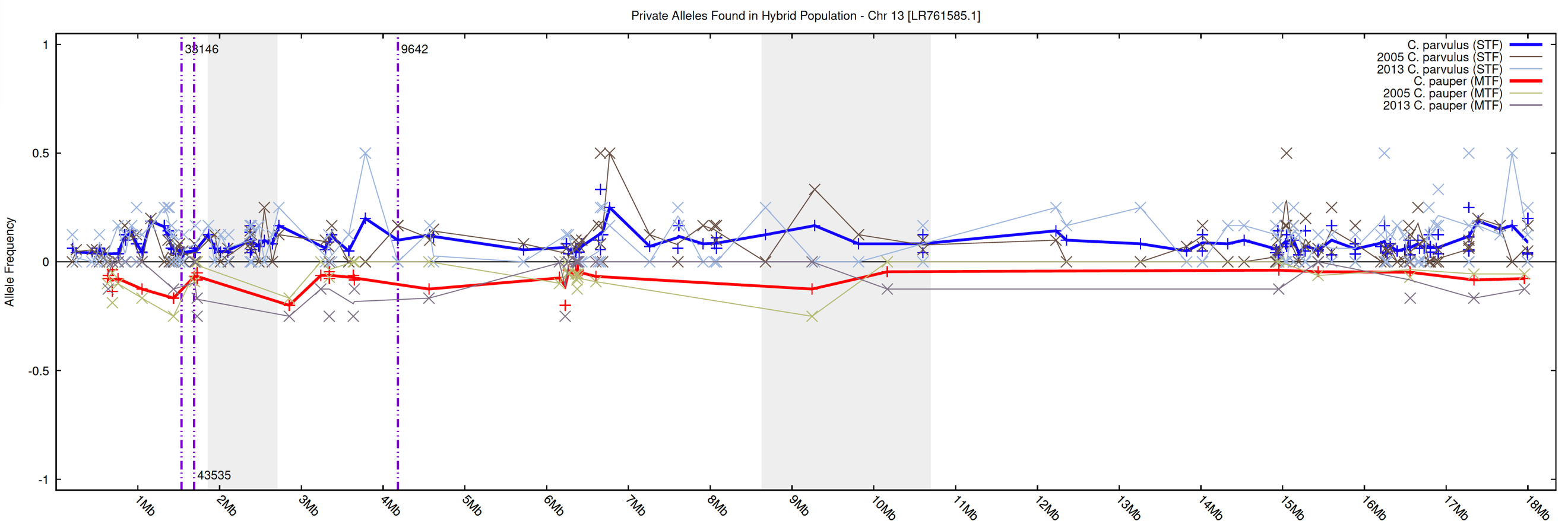

15

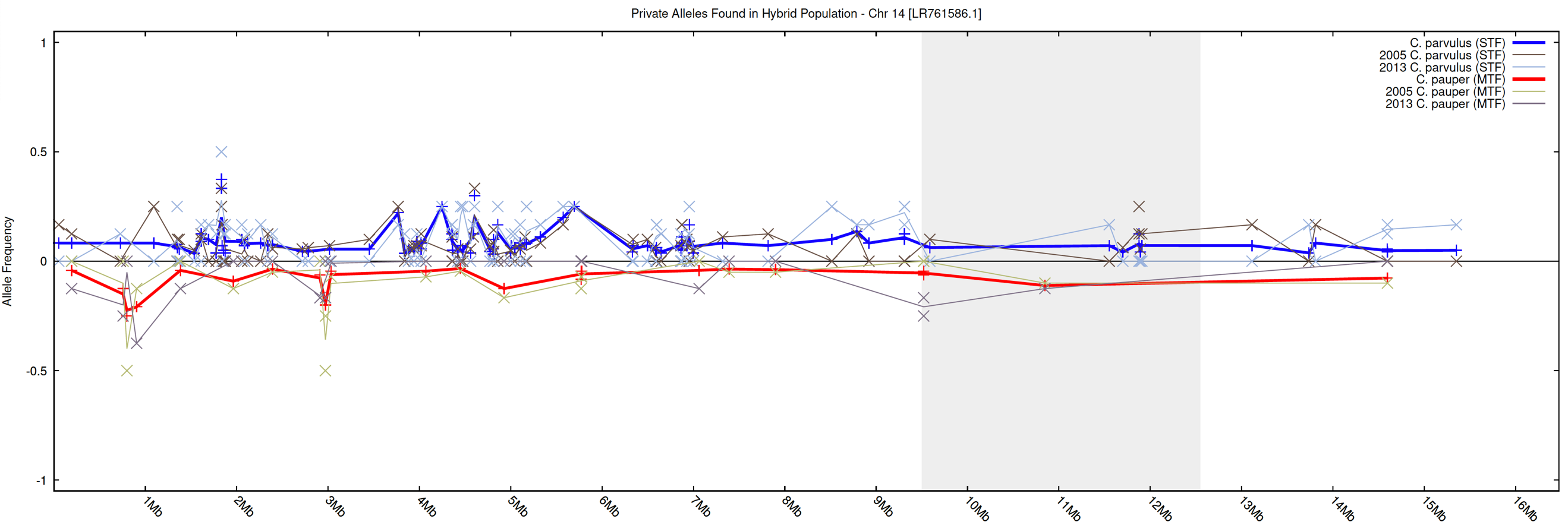

16

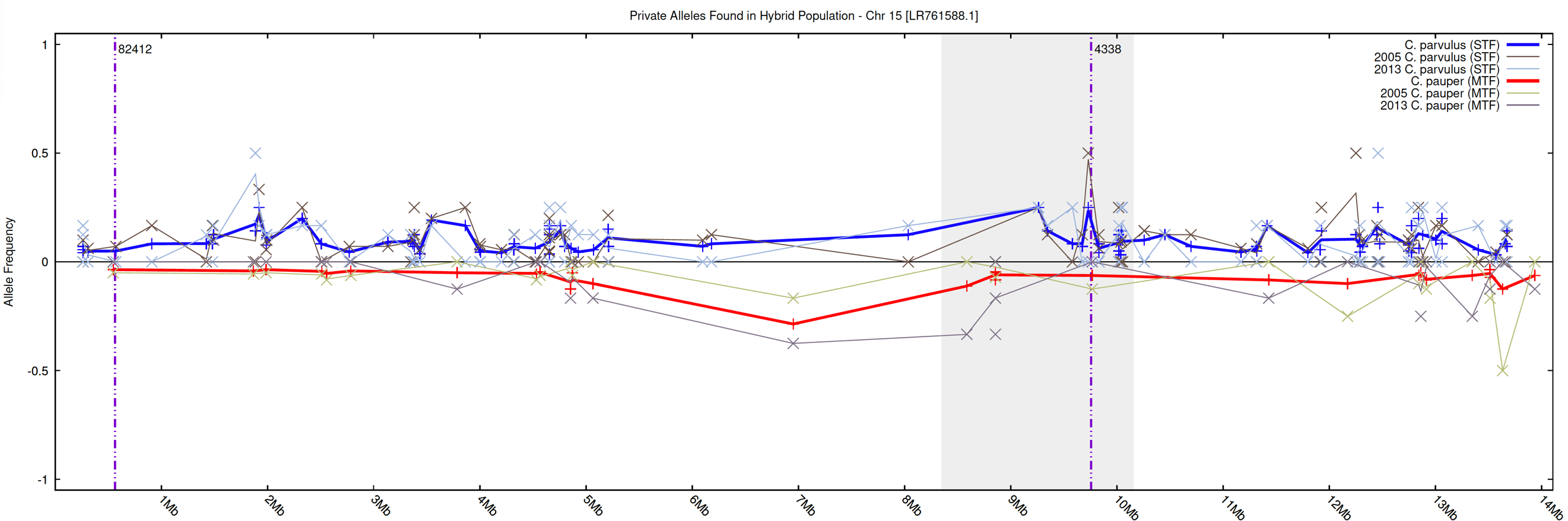

17

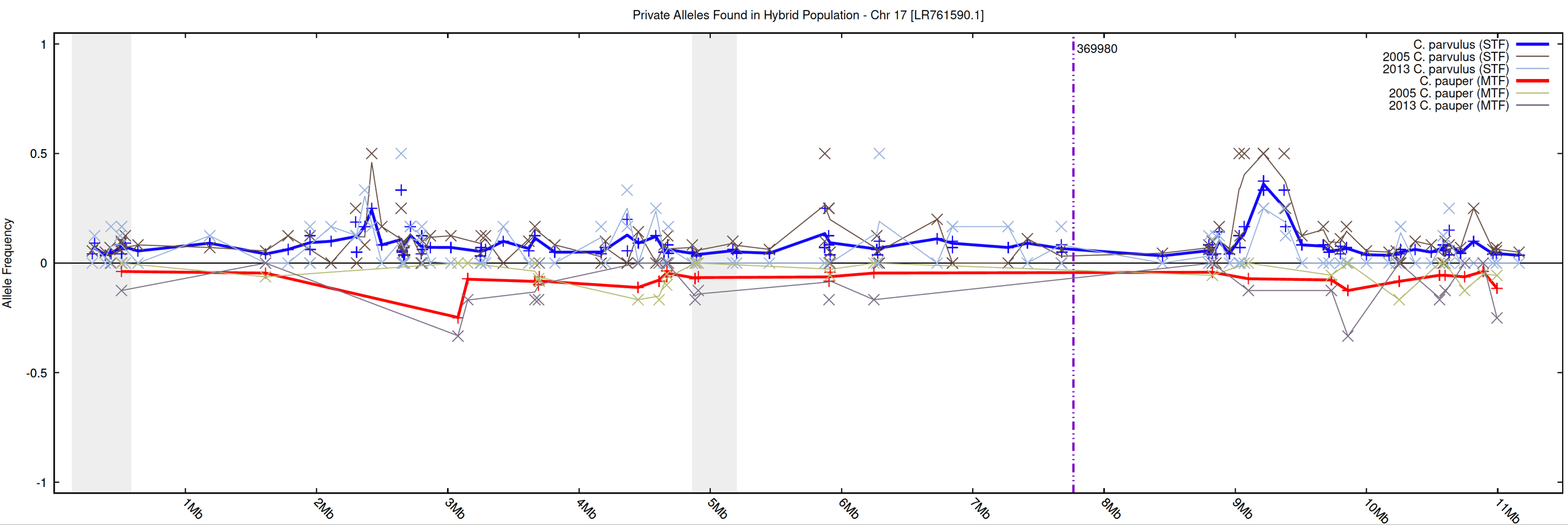

18

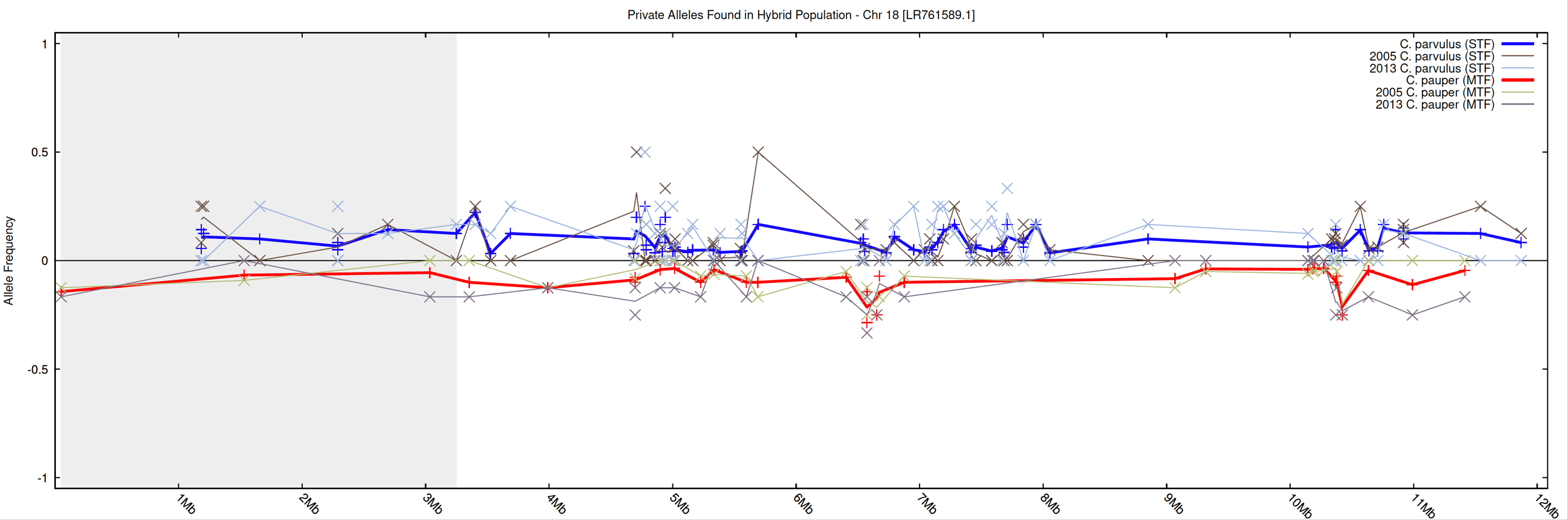

19

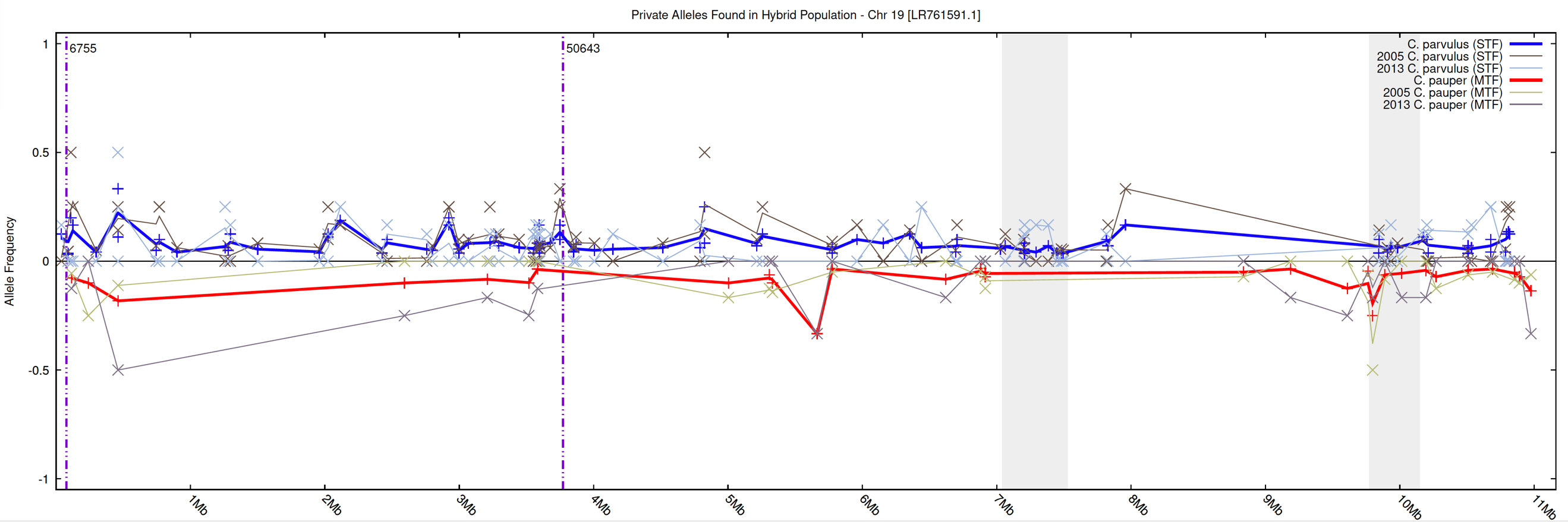

20

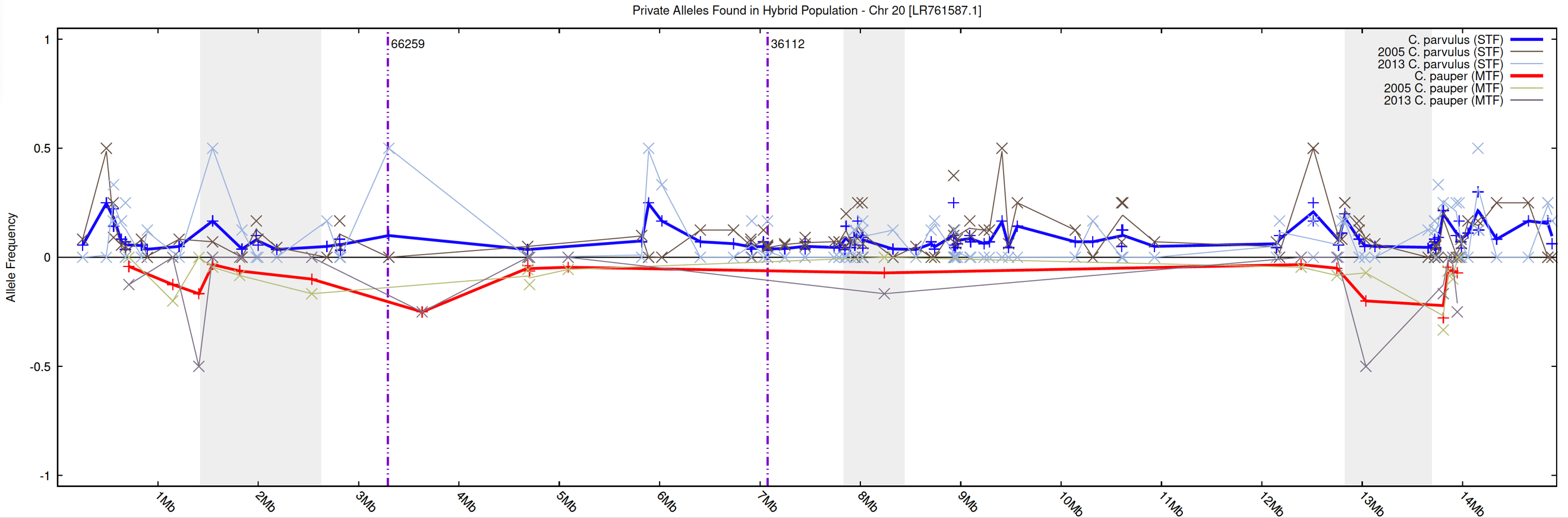

21

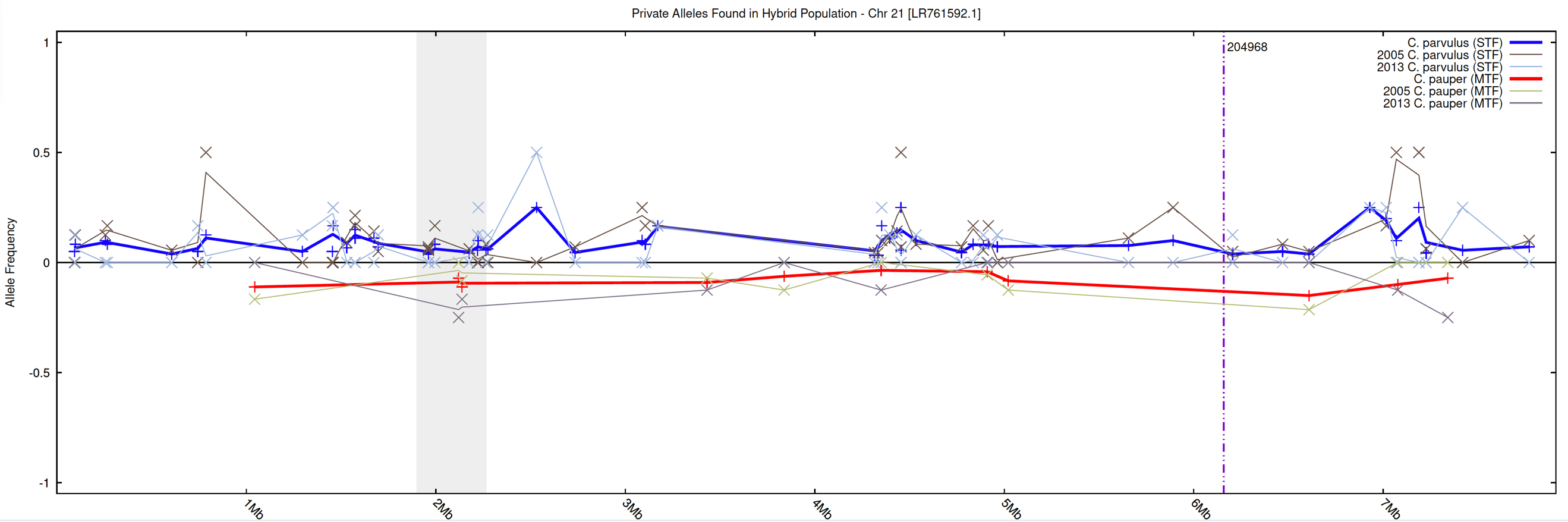

22

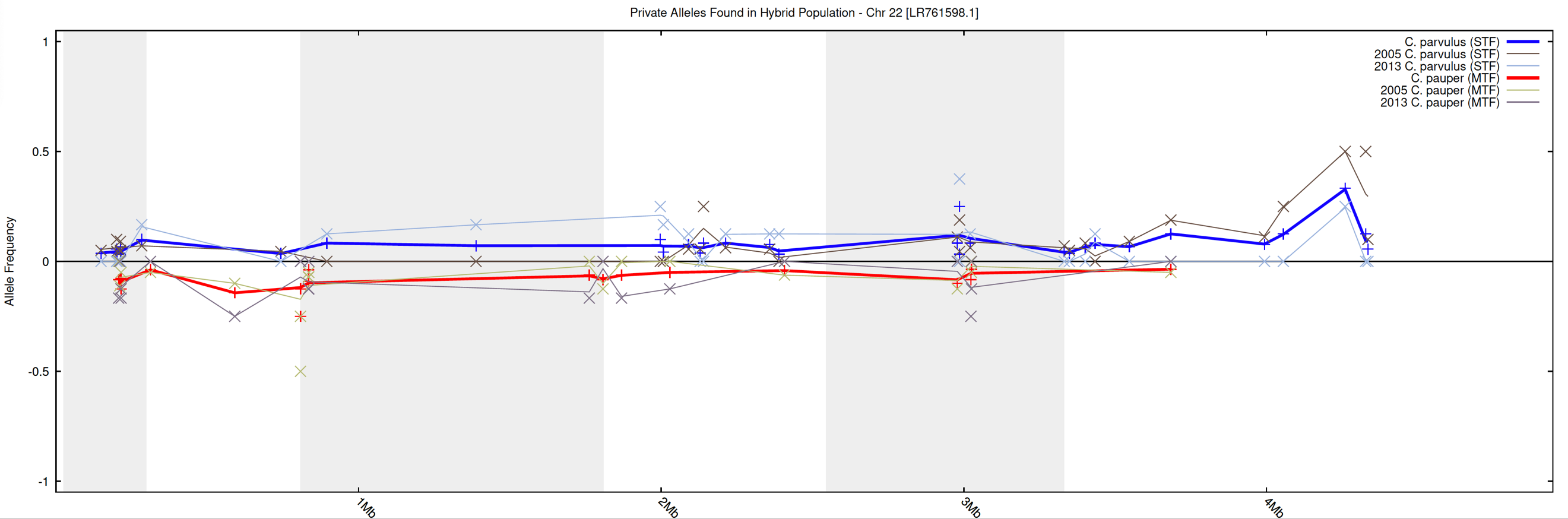

23

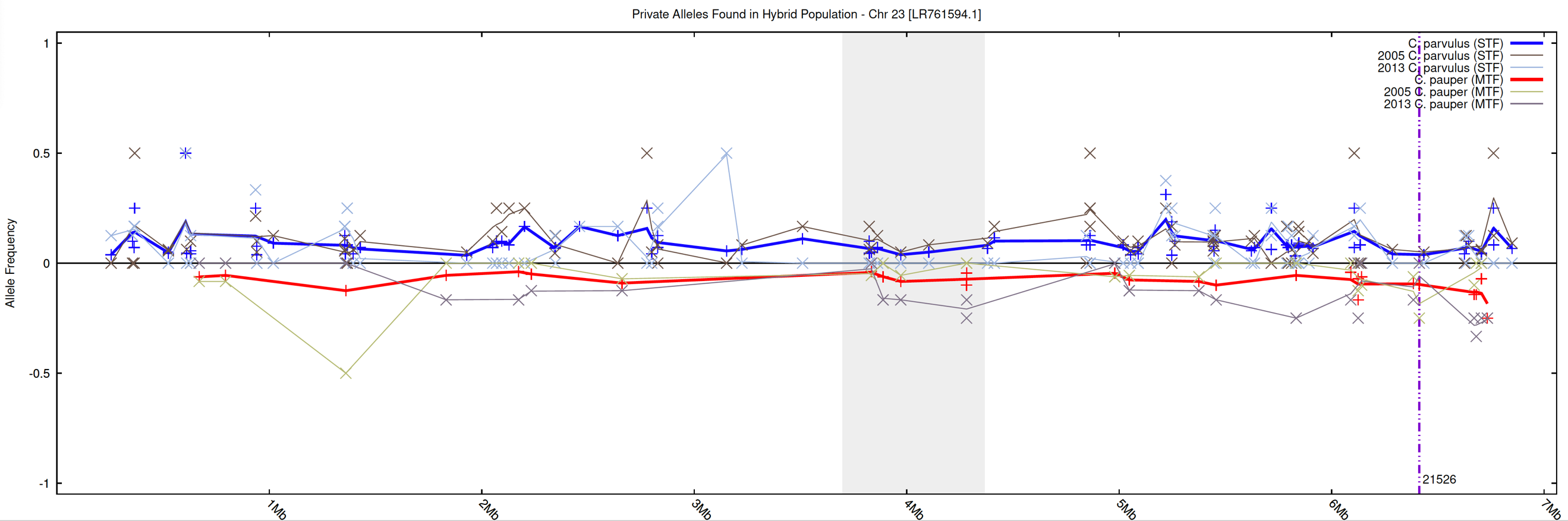

24

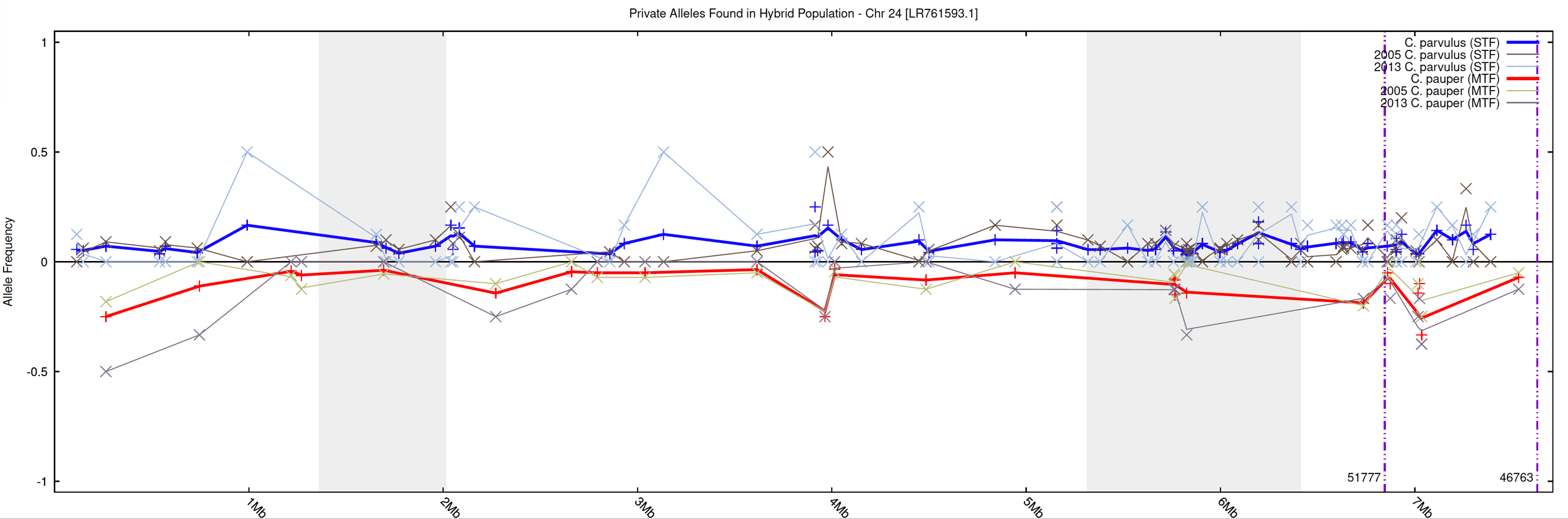

25

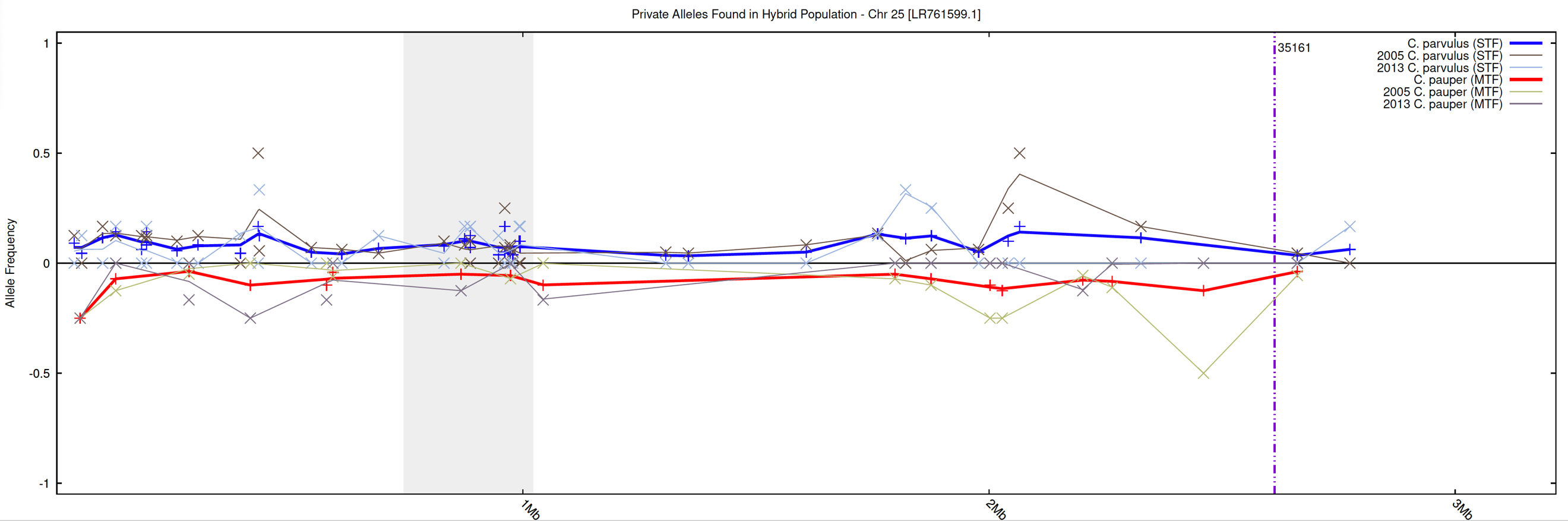

26

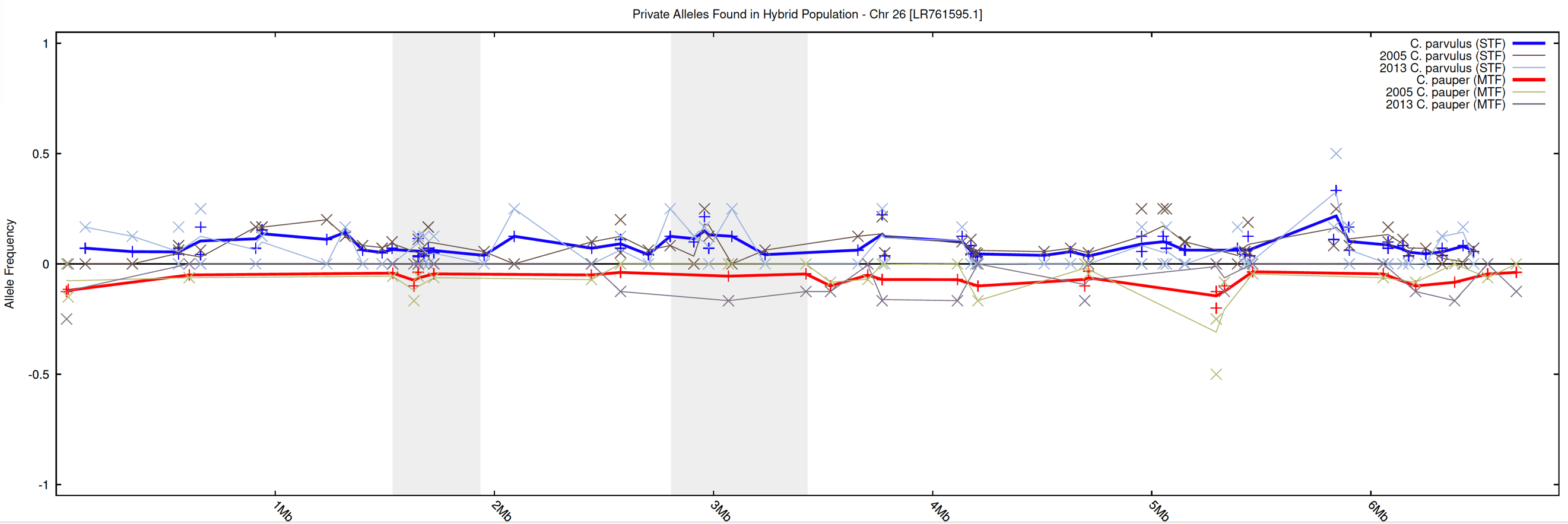

27

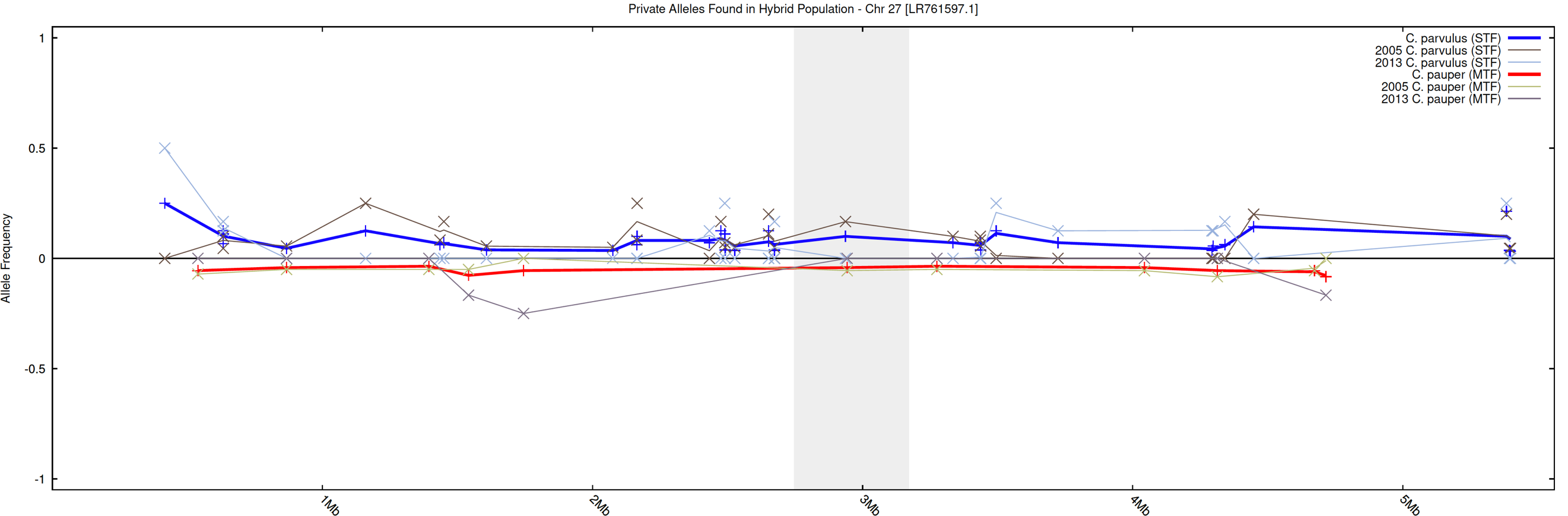

28

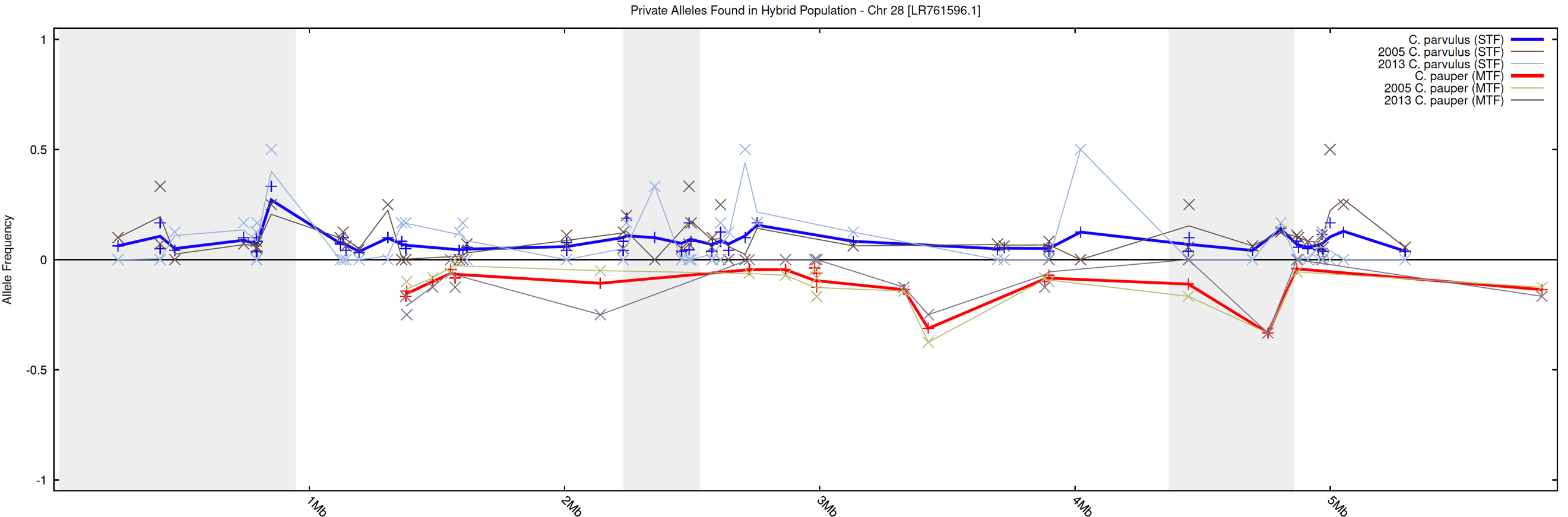

29

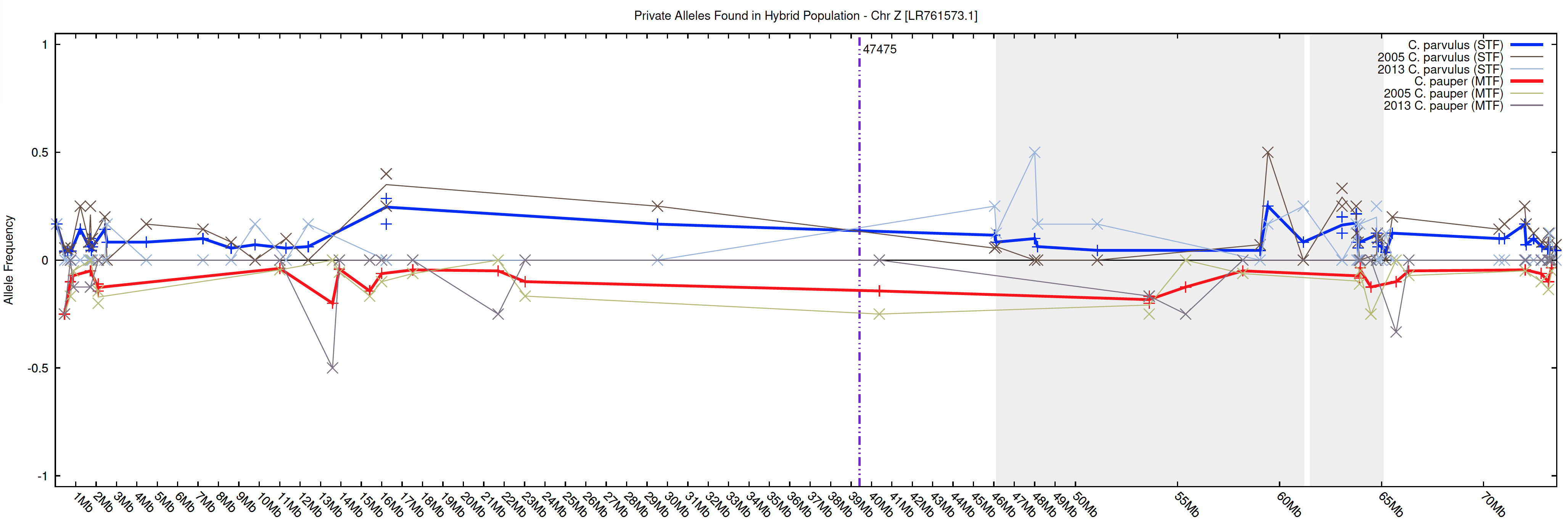

30

**Figure S7.** Private allele frequencies for hybrid C*amarhynchus* birds in 2005 versus 2013 mapped on to the *C. parvulus* chromosomes (1-30), coloured by parental species of origin (blue = *C. parvulus* and red = *C. pauper*). Dark red and blue lines indicate averages across both years. Light lines show allele frequency changes from 2005-2013. Shaded areas of the chromosome are regions where outlier windows were identified using local PCA. Horizontal dashed lines indicate the location of candidate loci under selection using *pcadapt*.
